# *Drosophila* Heat Shock Factor (HSF) Regulates Developmental Growth by Maintaining the Basal Expression of HSP83/HSP90

**DOI:** 10.64898/2026.04.17.719278

**Authors:** Jinghong J. Tang (唐璟鸿), Alicia Shipley, Roger P. White, Elijah Ferreira, Somes Schwinghammer, Michael A. Welte

## Abstract

The heat shock transcription factor HSF1 is best known as a master regulator of the proteotoxic stress response, yet its functions in animal development remain incompletely defined. In *Drosophila melanogaster*, heat shock factor (HSF) is essential for viability, but the mechanisms by which it promotes development are unclear. Here, we show that *Hsf* null larvae arrest at the early 2^nd^ instar stage and exhibit a significant reduction in basal levels of the chaperone HSP83/HSP90. Tissue-specific knockdown of *Hsf* revealed widespread and organ-specific requirements, including defects in endoreplication and cell growth in larval prothoracic and salivary glands, adult wing defects following larval imaginal disc perturbation, follicle degeneration in the ovary, and melanotic tumor upon hemocyte depletion. In these tissues, loss of HSF typically leads to reduced HSP83 levels, and restoration of HSP83 expression partially or fully rescues these defects. These findings identify HSP83 as a critical downstream effector of HSF and demonstrate that HSF promotes development largely by maintaining basal chaperone expression. Together, our results establish HSF as a key regulator of developmental progression and highlight a central role for proteostasis in supporting tissue growth under non-stress conditions.

## INTRODUCTION

Essentially all organisms respond to elevated temperatures by inducing a conserved stress response pathway, the heat shock response, that leads to the upregulation of a set of molecular chaperones. For example, in cultured *Drosophila* cells, the chaperone HSP70 is virtually undetectable at 25°C (Velazquez et al., 1983) but accumulates to become one of the most abundant proteins in the cell within a couple of hours after a shift to 37°C (Lindquist, 1980a, 1980b). Other chaperones, like *Drosophila* HSP83/HSP90, are already present in large amounts, but their levels rise even further after heat shock (Chomyn and Mitchell, 1982; Pauli et al., 1990; Riddihough and Pelham, 1986). Multiple mechanisms work together to allow for a rapid, yet massive response, including regulation of transcription, mRNA stability, translation, and post-translational mechanisms (McGarry and Lindquist, 1985; Theodorakis and Morimoto, 1987; Gomez-Pastor et al., 2018; Alagar Boopathy et al., 2022). Across eukaryotes, the transcriptional arm of the response is mediated by the transcription factor Heat Shock Factor (HSF1). HSF1 is activated in response to not only heat shock but many other proteotoxic types of stress, binds to target DNA sequences called Heat Shock Elements (HSEs) and induces the robust transcription of many chaperone genes (Mendillo et al., 2020). While the role of HSF1 in mediating the heat shock response has been characterized in great detail, its functions under non-stress conditions remain much less well understood. Notably, HSF1 is also critical for animal development in the absence of stress (Li et al., 2017).

Specific developmental functions for HSF1 have been explored in *Caenorhabditis elegans* and in mammals. In *C. elegans*, loss of HSF1 results in arrest at the late L2/early L3 stage, and, working together with the transcription factor E2F, HSF regulates a developmental transcriptional program that is distinct from the canonical stress response (Li et al., 2016). In mammals, HSF1 is similarly required for normal development, as mutant mice exhibit female infertility, postnatal growth retardation, and neurodevelopmental defects (Christians et al., 2000; Takaki et al., 2006; Xiao et al., 1999). Despite these organismal phenotypes, the tissue-specific requirements of HSF during development remain poorly defined. In *C. elegans*, germline-specific depletion of HSF impairs proliferation as well as early meiotic progression and is accompanied by downregulation of proteostasis-related genes, suggesting a role in germline homeostasis (Edwards et al., 2021). However, the specific mechanisms by which HSF promotes developmental progression remain largely unknown.

In *Drosophila melanogaster*, heat shock factor (here known as HSF) was identified over 35 years go (Clos et al., 1990), yet its developmental roles remain incompletely defined. HSF is essential for viability, as *Hsf* null mutants die in early larval stages, and it is also required in the female germline, where its loss causes follicle arrest during mid-oogenesis (Jedlicka et al., 1997). In cultured cells, ChIP–chip analysis identified HSF binding at ∼200 chromatin regions (Gonsalves et al., 2011), raising the possibility that HSF promotes the expression of many developmentally important genes. If so, its role could be similar to that of mammalian HSF1 in cancers, where it promotes cancer growth and drives a transcriptional program distinct from the canonical stress response (Dai et al., 2007; Mendillo et al., 2012; Santagata et al., 2013). However, the *in vivo* developmental roles of HSF in *Drosophila* remain unclear. In particular, it is unknown in which tissues HSF is required and which target genes are functionally important. While HSF has been shown to maintain basal expression of HSP83 in cultured cells and larval salivary glands, the extent to which this function generalizes across tissues is unknown (Gonsalves et al., 2011; Salamanca et al., 2011). In fact, earlier analyses using *Hsf* null clones and a temperature-sensitive allele (*Hsf^4^*) suggested that HSF plays only limited roles in viability and growth beyond early larval stages and in the female germline (Jedlicka et al., 1997). Thus, while HSF is clearly important for viability, both its key downstream effectors of HSF and the specific developmental stages at which it acts remain to be defined.

To identify how HSF promotes development, we examined its functions and downstream targets using classical alleles and tissue-specific knockdowns. We find that *Hsf* null larvae die at the early 2^nd^ instar stage, consistent with prior reports. In these larvae, basal HSP83 levels are dramatically reduced. Tissue-restricted knockdown revealed organ-specific requirements: reduced endoreplication and cell growth in the larval prothoracic and salivary glands, adult wing defects after larval imaginal-disc knockdown, degenerating follicles after knockdown in follicle cells, and melanotic masses upon hemocyte knockdown in larvae. Levels of HSP83 were reduced in affected tissues and in *Hsf* null larvae, and HSP83 re-expression partially or fully rescued the cellular and tissue phenotypes. These data indicate that HSF is essential for the development of numerous tissues and that this role depends to a large extent on maintaining basal HSP83 levels. Together, we identify HSF as a key regulator of development in *Drosophila* and define HSP83 as a biologically critical target.

## RESULTS

### HSF Is Essential for Early Larval Development

During larval development, *Drosophila* larvae increase their mass 200-fold within only three days, by tissue expansion through increases in both cell size and number (Church and Robertson, 1966; Nijhout, 2003; Thompson, 2010). Previous research suggests that *Hsf* is required for this massive growth as *Hsf* null mutants die in early larval stages (Jedlicka et al., 1997). To pinpoint the exact lethal phase, we analyzed two strong *Hsf* loss-of-function alleles. *Hsf^1^* is a nonsense mutation at residue 78, producing an unstable protein lacking all functional HSF domains (Jedlicka et al., 1997). *Hsf^03091^*is a loss-of-function mutation generated by P-element insertion (Ida et al., 2009; Spradling et al., 1999). We tracked developmental transitions (molting) in wild-type and trans-heterozygous *Hsf* mutants (*Hsf ^1/03091^*), as well as in animals where one of the two alleles was combined with a large deficiency that deletes *Hsf* and neighboring genes (*Hsf^1^/Df,* and *Hsf^03091^/Df*). At 24 hours after hatching (hAH), most wild-type larvae had molted into the 2^nd^ instar stage, whereas *Hsf* mutants remained in the 1^st^ instar stage (Fig. 1A, B). By 30 hAH, most *Hsf* mutants were still in the 1^st^ instar stage, with a few reaching the 2^nd^ instar stage (Fig. 1A, B). Shortly after molting, *Hsf* mutants died as 2^nd^ instar larvae, confirmed by their 2^nd^ instar mouth hooks (Fig. 1A, B, and C). A small proportion of 1^st^ instar *Hsf* null larvae survived up to 48 hAH before molting and subsequently dying as 2^nd^ instar larvae (Fig. 1B). Unlike wild-type larvae, which increased in size after molting, many 2^nd^ instar *Hsf* null larvae appeared smaller than their 1^st^ instar counterparts, indicating a growth defect (Fig. 1A). To confirm that these phenotypes are indeed the results of lack of *Hsf*, we also examined in the same experiments *Hsf ^1/03091^*mutants that also carry a tagged *Hsf* transgene under endogenous control (Tang et al., 2022); in this genotype, lethality and growth defects were rescued (Fig. 1A, B). We conclude that *Hsf* null mutants exhibit an extended 1^st^ instar stage and die shortly after molting into the 2^nd^ instar stage, suggesting that *Hsf* is essential for *Drosophila* larval developmental growth.

**Fig. 1.**
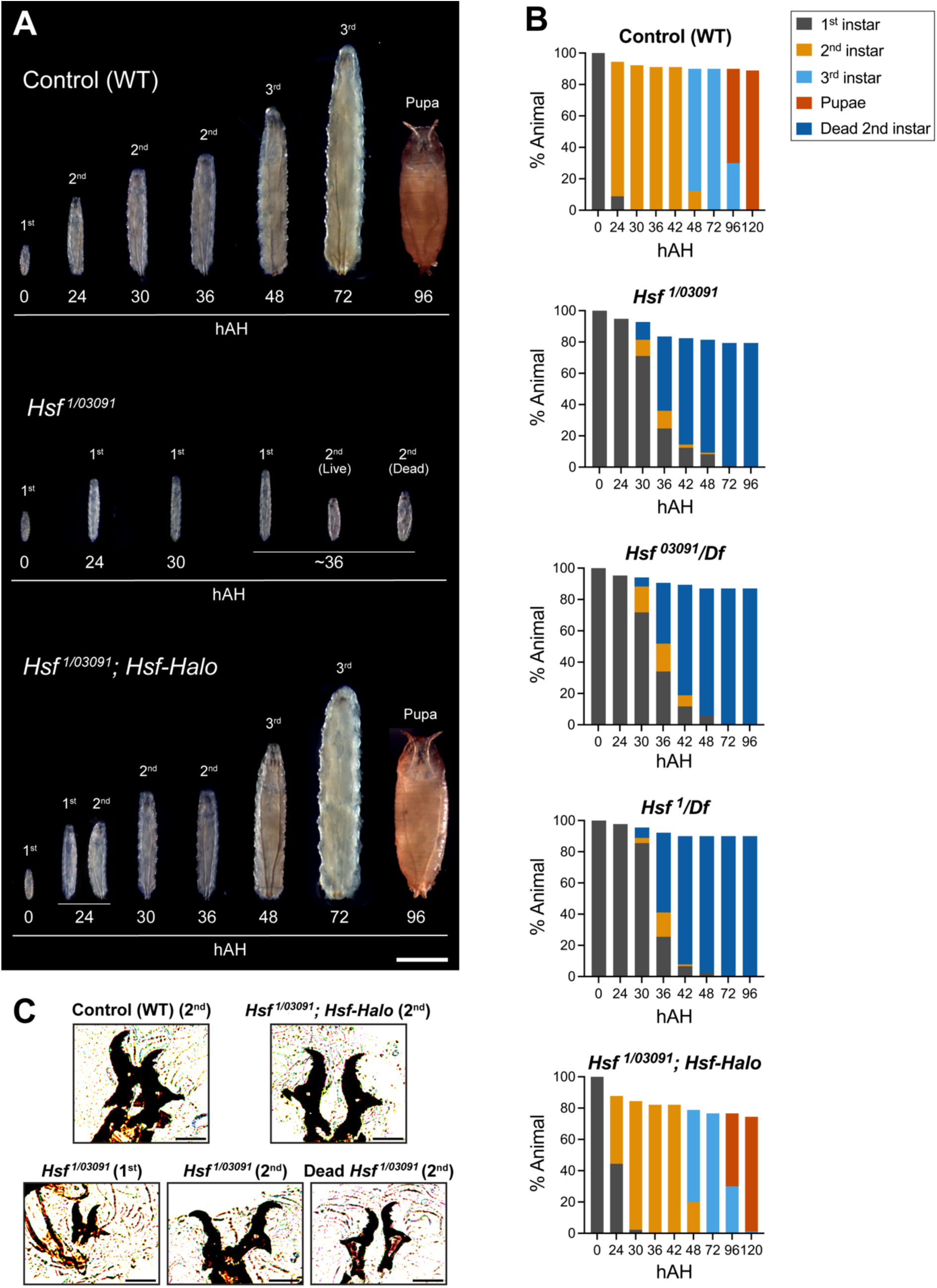
*Hsf* is required for early larval development. (A) Representative images of control (*Oregon R*), *Hsf* mutant (*Hsf^1/03091^*), and *Hsf* mutant larvae carrying the *Hsf-Halo* transgene (*Hsf^1/03091^; Hsf-Halo*) at different developmental stages. hAH, hours after hatching. Scale bar, 1mm. (B) Quantitation of developmental stage distribution and survival (%) in control (*Oregon R*), *Hsf* mutants (*Hsf^1/03091^*, *Hsf^03091^/Df*, and *Hsf^1^/Df*), and rescued animals (*Hsf^1/03091^; Hsf-Halo*). *Df* corresponds to *Df(2R)ED3610, a deficiency that encompasses the Hsf locus.* Color coding: 1^st^ instar larvae (grey), 2^nd^ instar larvae (orange), 3^rd^ instar larvae (light blue), and pupae (red). Dead 2^nd^ instar larvae are shown in dark blue. n = 80-100 larvae per genotype. (C) Larval mouth hook morphology of control (*Oregon R*) and rescued animals (*Hsf^1/03091^; Hsf-Halo*) at the 2^nd^ instar stage (upper panels). For *Hs*f mutants (*Hsf^1/03091^*), representative mouth hooks from 1^st^ instar, live 2^nd^ instar, and dead 2^nd^ instar larvae are shown (lower panels). Scale bar, 25 μm.

### HSP83 Is Down Regulated in *Hsf* Null Mutants

Heat-shock proteins (HSPs) are well-established targets of HSF and play central roles in maintaining proteostasis (Lindquist and Craig, 1988; Hu et al., 2022). Among developmentally expressed HSPs, Heat-shock Protein 83 (HSP83), the *Drosophila* ortholog of human HSP90, is expressed throughout development, including during the period of massive larval growth (Mason et al., 1984; Xiao and Lis, 1989). In the larval salivary gland, its basal expression requires HSF, consistent with ChIP-seq done in cultured *Drosophila* Kc cells showing strong HSF occupancy at the *Hsp83* promoter (Gonsalves et al., 2011; Salamanca et al., 2011).

To determine whether basal organismal HSP83 expression during larval development depends on HSF, we examined HSP83 protein levels using an *Hsp83-GFP* reporter (Tariq et al., 2009; Palumbo et al., 2020). We imaged HSP83–GFP–expressing larvae and quantified GFP across different developmental stages—newly hatched, late 1^st^ instar, and early 2^nd^ instar—in wild-type, *Hsf* null, and *Hsf* heterozygotes. We employed two metrics: normalized total GFP levels per larva and GFP density (GFP/area). For newly hatched larvae, *Hsf* null and *Hsf* heterozygous individuals showed slightly lower HSP83-GFP density but comparable overall levels of HSP83-GFP to wild-type (Fig. 2A, B, and C). However, for late 1^st^ instar and early 2^nd^ instar larvae, both HSP83-GFP levels and density were significantly reduced in *Hsf* null larvae compared to wild-type larvae, while *Hsf* heterozygous larvae were indistinguishable from wild type (Fig. 2A, B, and C). We also measured endogenous HSP83 levels in newly hatched (0 h AEH) and 24 h AEH wild-type and *Hsf* null larvae by Western blot. While newly hatched wild-type and *Hsf* null larvae had comparable HSP83 levels, *Hsf* null larvae showed significantly lower HSP83 levels at 24 h AEH compared to wild type (Fig. 2D, E). Thus, we conclude that HSF promotes normal HSP83 levels in the larva.

**Fig. 2.**
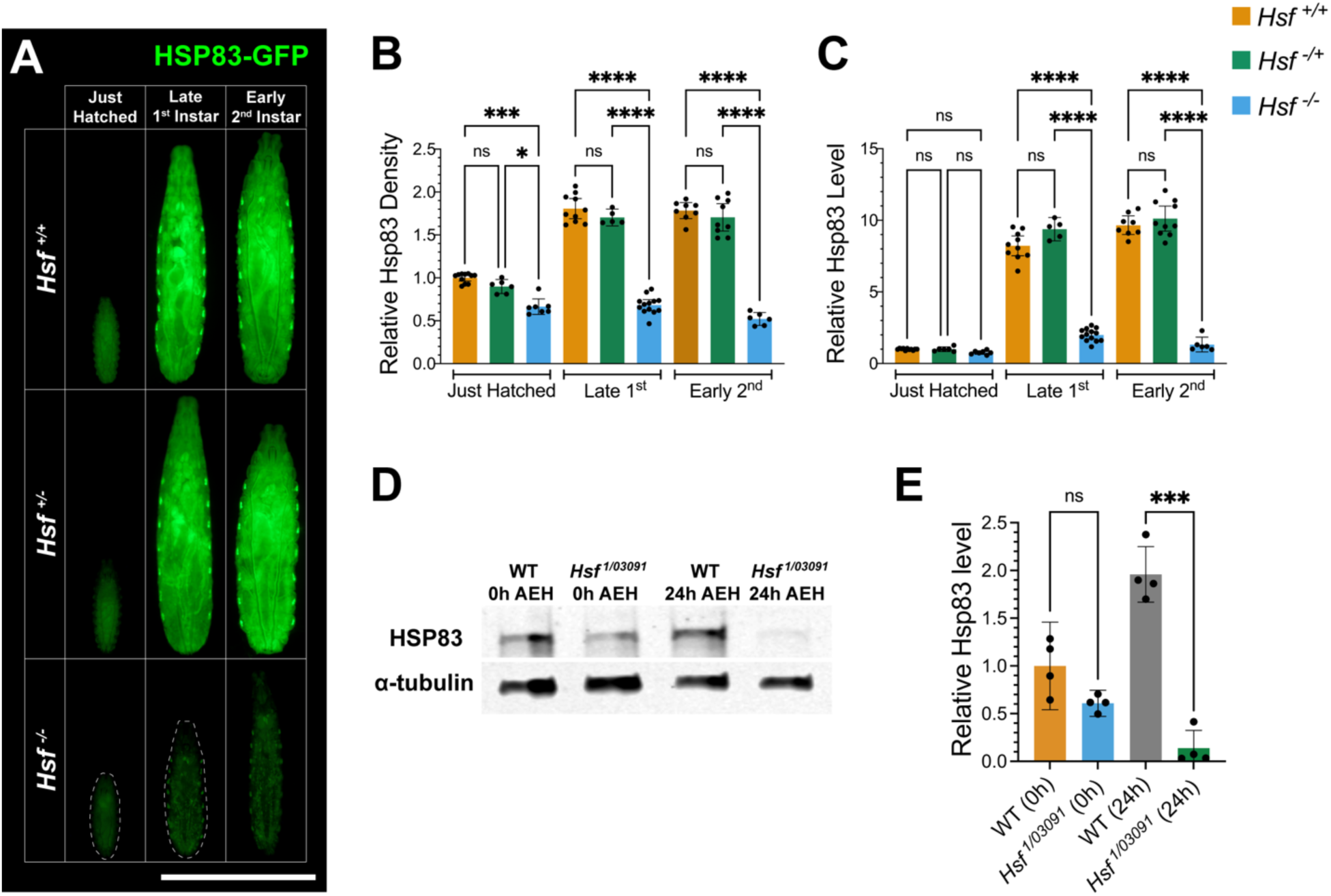
HSP83/HSP90 is downregulated in *Hsf* mutants. (A) Representative images of GFP fluorescence in HSP83–GFP-expressing control (*Hsf^+/+^*: *Oregon R*), *Hsf* heterozygous (*Hsf^+/-^*: *Hsf^1/+^* or *Hsf^03091/+^*) and *Hsf* mutant larvae (*Hsf^-/-^*: *Hsf^1/03091^*) at three developmental stages (newly hatched, late 1^st^ instar, and early 2^nd^ instar). GFP fluorescence is reduced in *Hsf* mutants. Scale bar, 1mm. (B) Quantitation of relative HSP83–GFP density (total HSP83 normalized to larval area) from (A). n = 5-13 larvae per genotype. (C) Quantitation of relative HSP83–GFP levels (total HSP83 per larva) from (A). n = 5-13 larvae per genotype. (D) Western blot analysis of endogenous HSP83 and α-tubulin in wild-type (WT) and *Hsf* mutant (*Hsf^1/03091^*) larvae at 0 and 24 h after egg hatching (AEH). (E) Quantitation of relative endogenous HSP83 levels from (D). n = 4 western blots. Statistical significance was determined using Brown-Forsythe and Welch ANOVA followed by Dunnett’s multiple comparisons test. Data are presented as mean ± 95% CI. P < 0.05 (*), P < 0.01 (**), P < 0.001 (***), P < 0.0001 (****).

### HSF Is Required for the Growth of the Prothoracic Gland

Given the early larval growth arrest of *Hsf* null mutants and the observed downregulation of HSP83 in these larvae, we hypothesized that HSF supports tissue growth by maintaining developmental proteostasis through regulating basal HSP83 expression. To test this hypothesis, we performed tissue-specific knockdown of *Hsf* using RNA interference in the rapidly growing larval prothoracic gland (PG), a secretory tissue within the larval ring gland, using two distinct hairpins driven by the PG-specific driver *phm-Gal4* (Fig. 3A, B). For subsequent imaging experiments, we also drove the expression of the membrane marker CD8-RFP, to be able to unambiguously identify the extent of the PG. Larvae with PG-specific *Hsf* knockdown failed to initiate pupariation, arresting at the 3^rd^ instar stage (Fig. 3B). Instead, they continued foraging for days and grew in size, exhibiting a giant larva phenotype (Fig. S1A, B). One important function of the PG is to produce the steroid hormone ecdysone, which triggers molting and metamorphosis (Ou et al., 2016; Yamanaka et al., 2013) (Fig. 3A). Feeding 20E to the *Hsf* knockdown larvae rescued the developmental arrest and induced pupariation, suggesting that *Hsf* is essential for normal levels of ecdysone production in the PG (Fig. S1C, D).

**Fig. 3.**
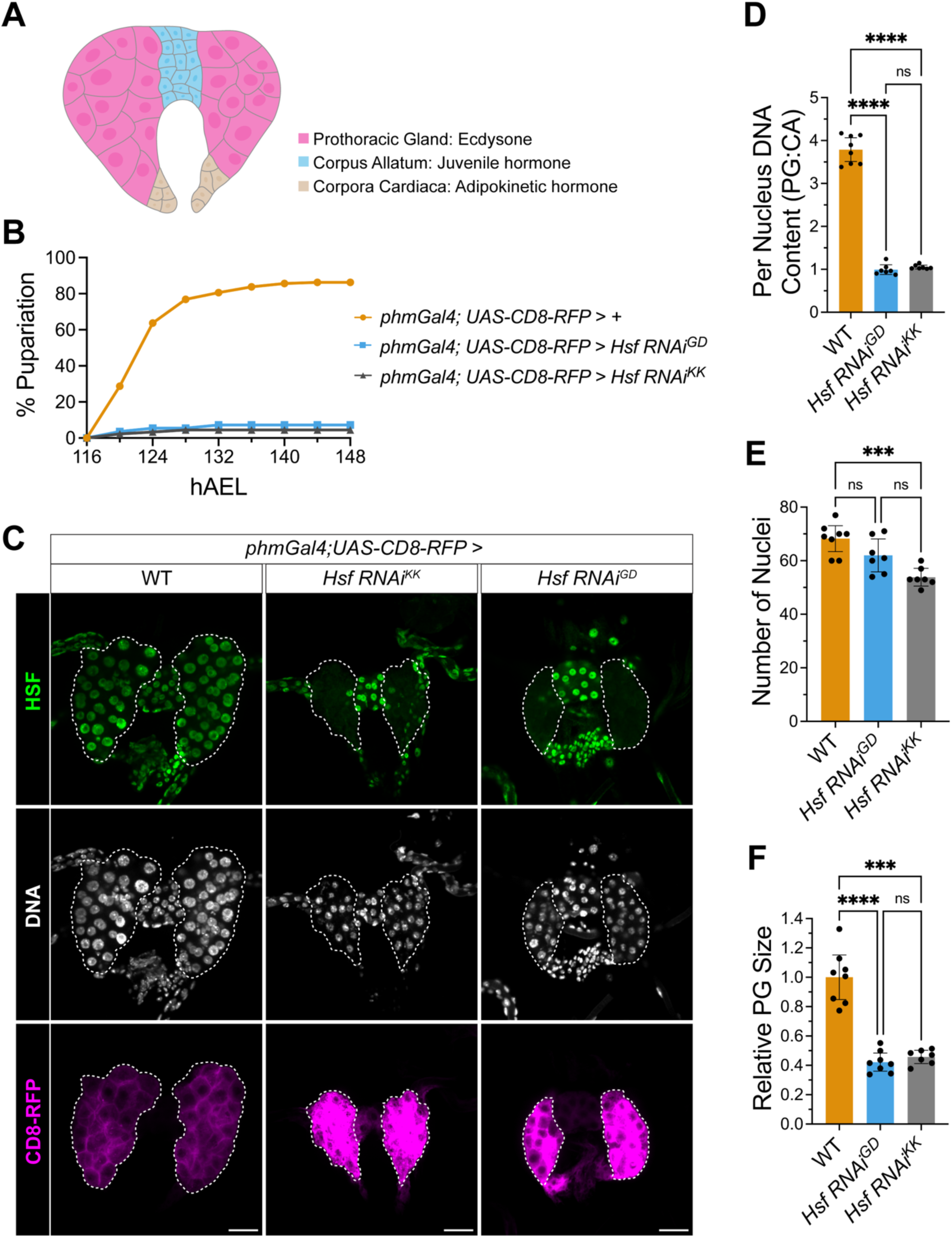
Prothoracic gland (PG)-specific knockdown of *Hsf*. (A) Graphic illustration of the *Drosophila* ring gland. Pink: Prothoracic gland (PG) that synthesizes ecdysone. Blue: Corpus Allatum (CA) that synthesizes juvenile hormone. Brown: Corpora Cardiaca that synthesizes adipokinetic hormone. (B) Percent pupariation over time in control and two independent PG-specific *Hsf* knockdown lines (*Hsf RNAi^GD^*, and *Hsf RNAi^KK^*) from 116 to 148 h after egg laying (hAEL). n = 160, 110, and 90 for control, *Hsf RNAi^GD^*, and *Hsf RNAi^KK^*, respectively. (C) Immunostaining for HSF (green), Hoechst staining for DNA (white), and CD8-RFP expression (magenta) in the prothoracic gland of wandering 3^rd^ instar larvae at ∼120 h AEL in control and *Hsf RNAi* animals. The prothoracic gland is outlined with white dashed lines. Scale bar, 20 μm. (D) Quantitation of DNA content per PG nucleus, normalized to corpus allatum (CA) nuclei, in control and *Hsf RNAi* PGs from (C). n = 7-8 PGs per genotype. (E) Quantitation of the number of nuclei per PG in control and *Hsf RNAi* animals from (C). n = 7-8 PGs per genotype. (F) Quantitation of relative PG size in control and *Hsf RNAi* animals from (C). n = 7-8 PGs per genotype. Statistical significance was determined using Brown-Forsythe and Welch ANOVA followed by Dunnett’s multiple comparisons test. Data are presented as mean ± 95% CI. P < 0.05 (*), P < 0.01 (**), P < 0.001 (***), P < 0.0001 (****).

During normal development, the PG undergoes massive growth, initially via mitoses during the 1^st^ instar stage and then via endoreplication during the 2^nd^ and 3^rd^ instar stages, with individual cells reaching a DNA content of 64C (Ohhara et al., 2017). To determine whether *Hsf* deficiency affects PG growth, we dissected PGs and performed HSF and DNA staining. HSF immunostaining showed that HSF is localized in the nuclei (Fig. 3C), consistent with previous findings that HSF is predominantly nuclear in *Drosophila* cells (Wang and Lindquist, 1998). *Hsf* RNAi PGs had little to no HSF signal, were smaller, and had reduced DNA content (Fig. 3C, D, and F). One hairpin (KK) caused a slight decrease in cell number, while the other (GD) had no effect. These findings suggest that HSF is essential for prothoracic gland growth and endoreplication.

### HSP83 Is Downregulated in *Hsf* RNAi PG and Necessary for PG Growth

We next tested whether HSF is required for normal expression of HSP83, like in whole larvae. We imaged PGs from control and *Hsf* RNAi larvae expressing the HSP83–GFP reporter and labeled PGs by staining for disembodied (Dib), a cytochrome P450 involved in ecdysteroid biosynthesis and specifically expressed in the PG (Parvy et al., 2005). We then quantified two metrics: normalized total GFP levels per gland and GFP density (GFP/area). We found that both HSP83-GFP levels and density were significantly reduced in *Hsf* RNAi PGs (Fig. 4A, C, and D). Because these glands fail to grow, it is conceivable that these reduced levels are a secondary effect of reduced PG size. We therefore aimed to reduce gland size via an independent genetic manipulation. We knocked down the DNA replication-related element-binding factor (Dref). *Dref* lacks predicted heat shock factor binding elements (HSEs) and shows no HSF binding by ChIP-seq (Gonsalves et al., 2011). *Dref* RNAi resulted in phenotypes similar to *Hsf* knockdown: developmental arrest at the 3rd instar stage (Fig. S2A) and smaller PG size (Fig. 4A, B) (Park et al., 2012). Although HSP83-GFP level and density were lower in *Dref* RNAi PGs compared to wild type, they remained significantly higher than in *Hsf RNAi* PGs. Thus, the reduction in HSP83-GFP in *Hsf* RNAi PGs is not explained solely by organ size and supports a direct requirement for HSF in maintaining basal HSP83 expression in the PG.

**Fig. 4.**
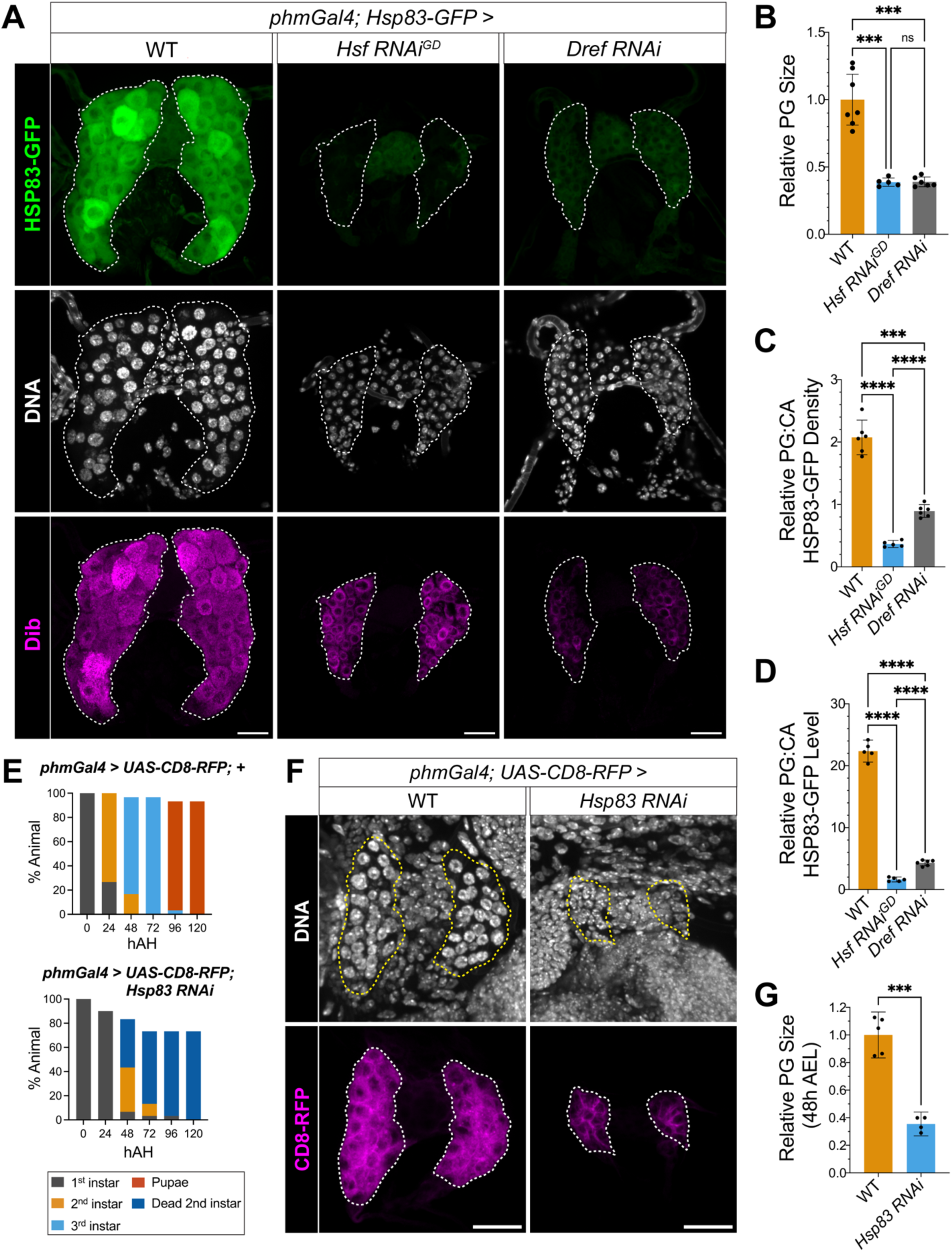
HSP83 is downregulated in *Hsf* knockdown prothoracic glands (PGs). (A) HSP83–GFP (green), Hoechst staining for DNA (white), and anti-Dib immunostaining (magenta) in the PGs of wandering 3^rd^ instar larvae at ∼120 h after egg laying (AEL) in control and PG-specific *Hsf* or *Dref* knockdown (*Hsf RNAi^GD^* and *Dref RNAi)* animals. The PG is outlined with white dashed lines. Scale bar, 20 μm. (B) Quantitation of relative PG size in control, *Hsf RNAi^GD^*, and *Dref RNAi* animals from (A). n = 5-7 PGs per genotype. (C) Quantitation of relative HSP83–GFP density (total PG signal normalized to PG area and then to the corpus allatum (CA)) from (A). n = 5-7 PGs per genotype. (D) Quantitation of relative HSP83–GFP level (total PG signal normalized to the CA) from (A). n = 5-7 PGs per genotype. (E) Quantitation of developmental stage distribution and survival (%) in control and PG-specific *Hsp83* knockdown larvae. n = 30 larvae per genotype. (F) Hoechst staining for DNA (white) and CD8-RFP expression (magenta) in the PG of 2^nd^ instar larvae at ∼48 h AEL in control and *Hsp83 RNAi* animals. The PG is outlined with white or yellow dashed lines. Scale bar, 20 μm. Statistical significance in (B), (C), and (D) was determined using Brown-Forsythe and Welch ANOVA followed by Dunnett’s multiple comparisons test. Statistical significance in (E) was determined using Welch’s t test. Data are presented as mean ± 95% CI. P < 0.05 (*), P < 0.01 (**), P < 0.001 (***), P < 0.0001 (****).

If the phenotypes in *Hsf* RNAi PG are due to reduced HSP83 levels, directly knocking down *Hsp83* should produce similar or even more severe effects. We therefore performed RNAi-mediated knockdown of *Hsp83* in the PG. Unlike *Hsf* RNAi larvae, which arrested at the 3^rd^ instar stage, *Hsp83* RNAi larvae exhibited developmental delays and died at the 2^nd^ instar stage (Fig. 4E and Fig. S2B). Feeding these larvae 20E partially rescued the phenotype, allowing them to progress to and arrest at the 3^rd^ instar stage (Fig. S2B, C). The PGs of the 2^nd^ instar *Hsp83* RNAi larvae were smaller than those of wild-type larvae (Fig. 4F, G). We also observed that 2^nd^ instar *Hsp83* RNAi larvae frequently wandered out of the food, unlike 2^nd^ instar wild-type larvae, which remained within it (Fig. S2C). This phenotype has been linked to defects in ecdysone signaling (Kugler et al., 2011). To further evaluate the potential defects in foraging behavior, we placed a food patch at the center of a 6 cm plate and recorded the time it took larvae, placed at the edge, to enter the food within one hour. While all wild-type larvae entered the food within 20 minutes, only about 30% of *Hsp83* RNAi larvae did so within one hour, indicating impaired food-seeking behavior (Fig. S2E). Feeding 20E to *Hsp83* RNAi larvae rescued the behavior, with all larvae entering the food within one hour (Fig. S2E). We conclude that HSP83 is required for PG growth and function, and its expression in the PG requires HSF.

### Restorating HSP83 Expression Rescues *Hsf* RNAi PG Phenotypes

Knockdown of *Hsf* or *Hsp83* in the PG result in similar phenotypes, and *Hsf* is necessary for normal expression levels for HSP83 in the PG. These observations suggest that the phenotypes observed with *Hsf* knockdown are largely caused by the loss of HSP expression. However, since *Hsf* has many potential targets (Gonsalves et al., 2011), *Hsf* knockdown might disrupt numerous additional pathways important for normal PG development. We therefore re-expressed HSP83 in the *Hsf* RNAi PGs using *UAS-Hsp83*. Restoring HSP83 expression did not affect the number of PG nuclei but fully restored DNA content to wild-type levels and partially rescued PG size (Fig. 5A, B, C, and D). Additionally, it partially rescued the developmental arrest of *Hsf* RNAi larvae: although the onset of pupariation was delayed, the pupariation rate matched that of wild-type larvae and those pupae eclosed (Fig. 5F). Overexpressing HSP83 alone in wild-type PGs caused only a slight increase in PG size without affecting the number of nuclei or DNA content, indicating that HSP83 overexpression by itself does not dramatically alter these properties (Fig. S3A, B, C, and D). These results suggest that *Hsf* regulates prothoracic gland endoreplication and growth by controlling the expression of HSP83.

**Fig. 5.**
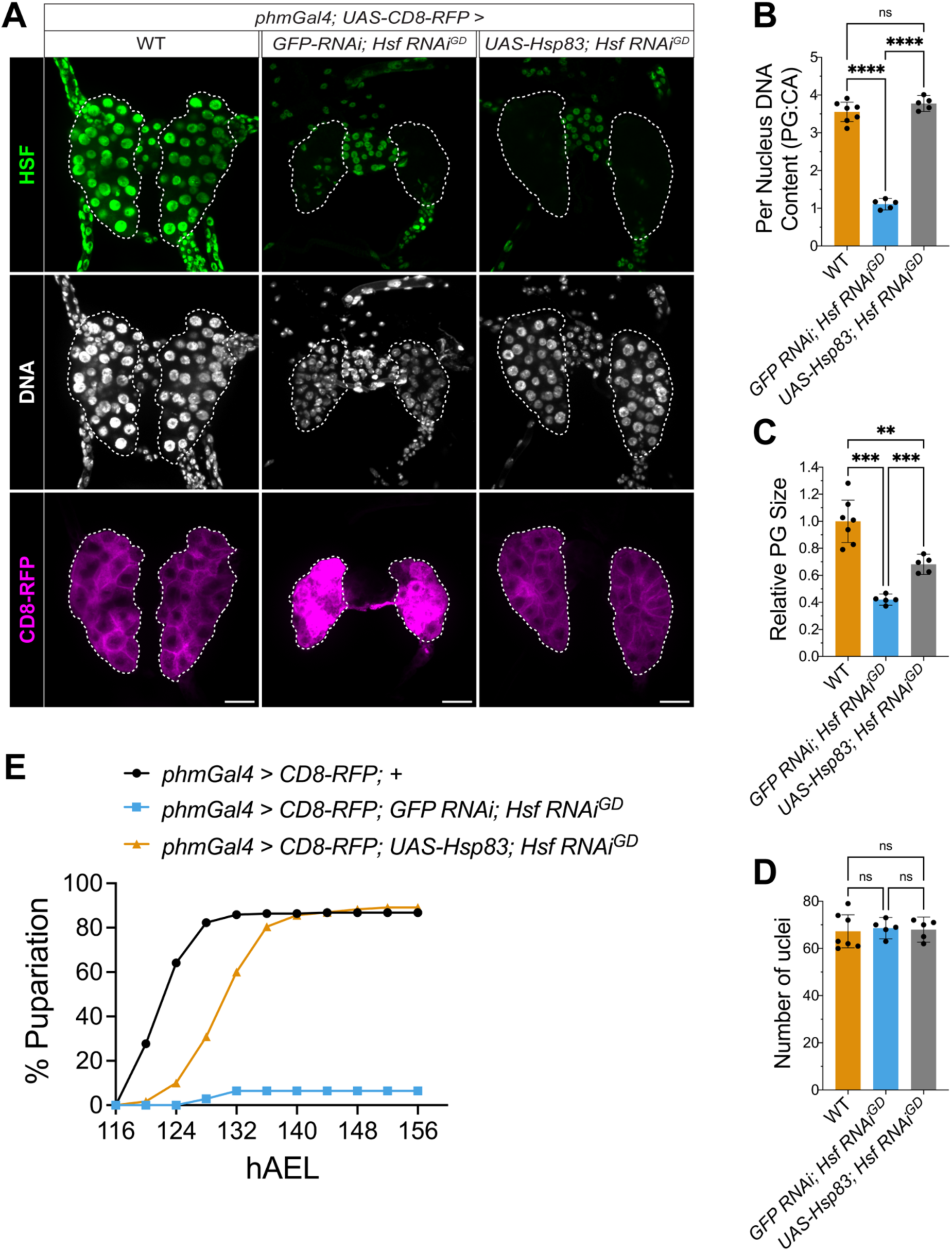
Restoration of HSP83 expression partially rescues prothoracic gland (PG)–specific *Hsf* knockdown phenotypes. (A) Immunostaining for HSF (green), Hoechst staining for DNA (white), and CD8-RFP expression (magenta) in the PG of wandering 3^rd^ instar larvae at ∼120 h after egg laying (hAEL) in control and *Hsf RNAi* animals, with or without HSP83 re-expression (*UAS-Hsp83*). The prothoracic gland is outlined with white dashed lines. Scale bar, 20 μm. (B) Quantitation of DNA content per PG nucleus, normalized to corpus allatum (CA) nuclei, in control and *Hsf RNAi* PGs, with or without HSP83 re-expression, from (A). n = 5-7 PGs per genotype. (C) Quantitation of relative PG size, in control and *Hsf RNAi* PGs, with or without HSP83 re-expression, from (A). n = 5-7 PGs per genotype. (D) Quantitation of the number of nuclei per PG, in control and *Hsf RNAi* PGs, with or without HSP83 re-expression, from (A). n = 7-8 PGs per genotype. (E) Percent pupariation over time in control and *Hsf RNAi* PGs, with or without HSP83 re-expression, from 116 to 156 h after egg laying (hAEL). n = 220, 140, and 230 for control, *Hsf RNAi* PGs, and *Hsf RNAi* PGs with HSP83 re-expression, respectively. Statistical significance was determined using Brown-Forsythe and Welch ANOVA followed by Dunnett’s multiple comparisons test. Data are presented as mean ± 95% CI. P < 0.05 (*), P < 0.01 (**), P < 0.001 (***), P < 0.0001 (****).

As the master regulator of the heat-shock response, HSF is known to upregulate HSP83 under stress conditions, raising the possibility that the HSF-dependent HSP83 expression we observe in the prothoracic gland (PG) reflects developmental stress. Because HSP70 is negligible at baseline and robustly induced by HSF during stress (Velazquez et al., 1983), we used HSP70 as a stress readout. A 20-min heat shock induced strong HSP70 signal in the PG detected by immunostaining, whereas HSP70 was undetectable in PGs from untreated wild-type larvae (Fig. S4A and B). Thus, HSF’s role during PG growth is unlikely to reflect activation of the canonical heat-shock program.

### HSF Is Essential for Salivary Gland Growth by Regulating HSP83

We next asked whether the requirement for HSF is specific to the prothoracic gland or extends to other developing tissues. Endoreplication is a common mechanism that regulates cell size without increasing cell number (Shu et al., 2018). Many *Drosophila* tissues grow through endoreplication, including adult nurse cells, adult follicle cells, fat bodies, and larval salivary glands (Smith and Orr-Weaver, 1991; Edgar and Orr-Weaver, 2001). To study whether *Hsf* also regulates endoreplication through HSP83 in other tissues, we knocked down *Hsf* in the larval salivary gland (SG) using the salivary gland-specific drivers *AB1-Gal4* and *ptc-Gal4*. In wild-type SGs, HSF exhibited nuclear localization as detected by immunostaining; this nuclear signal was absent in *Hsf* RNAi glands (Fig. 6A). There was diffuse cytoplasmic signal, possibly a result of non-specific antibody staining due to changes in tissue properties after *Hsf* knockdown; in any case, the absence of nuclear signal suggests that HSF’s ability as a transcription factor should be drastically impaired. *Hsf* RNAi significantly reduced SG cell size and DNA content (Fig. 6A, C, and D). HSP83-GFP density and level were also reduced in the *Hsf* RNAi SGs (Fig. 6E, G, and H). Similar to the prothoracic gland, knocking down *Dref* resulted in smaller SGs, enabling us to test whether the reduction in HSP83-GFP was solely due to reduced gland size (Fig. 6E, F). In *Dref* RNAi SGs, HSP83-GFP levels were lower than in the wild type but significantly higher than in *Hsf* RNAi SGs, and HSP83-GFP density was even higher than the wild type (Fig. 6E, H). These findings indicate that HSP83 is specifically downregulated in *Hsf* RNAi SGs. Furthermore, knocking down *Hsp83* alone also reduced SG size and DNA content (Fig. 6B, C, and D). In addition, restoring HSP83 expression partially rescued the SG size and DNA content defects caused by *Hsf* RNAi (Fig. 6A, C, and D). Finally, we also knocked down *Hsf* in endoreplicating follicle cells using *traffic-jam-Gal4*, which resulted in degenerating follicles, as indicated by condensed DNA staining and the absence of late-stage follicles. This phenotype was partially rescued by re-expression of HSP83 (Fig. S5). Together, these results suggest that *Hsf* regulates endoreplication through HSP83 in many, and possibly all, endoreplicating tissues.

**Fig. 6.**
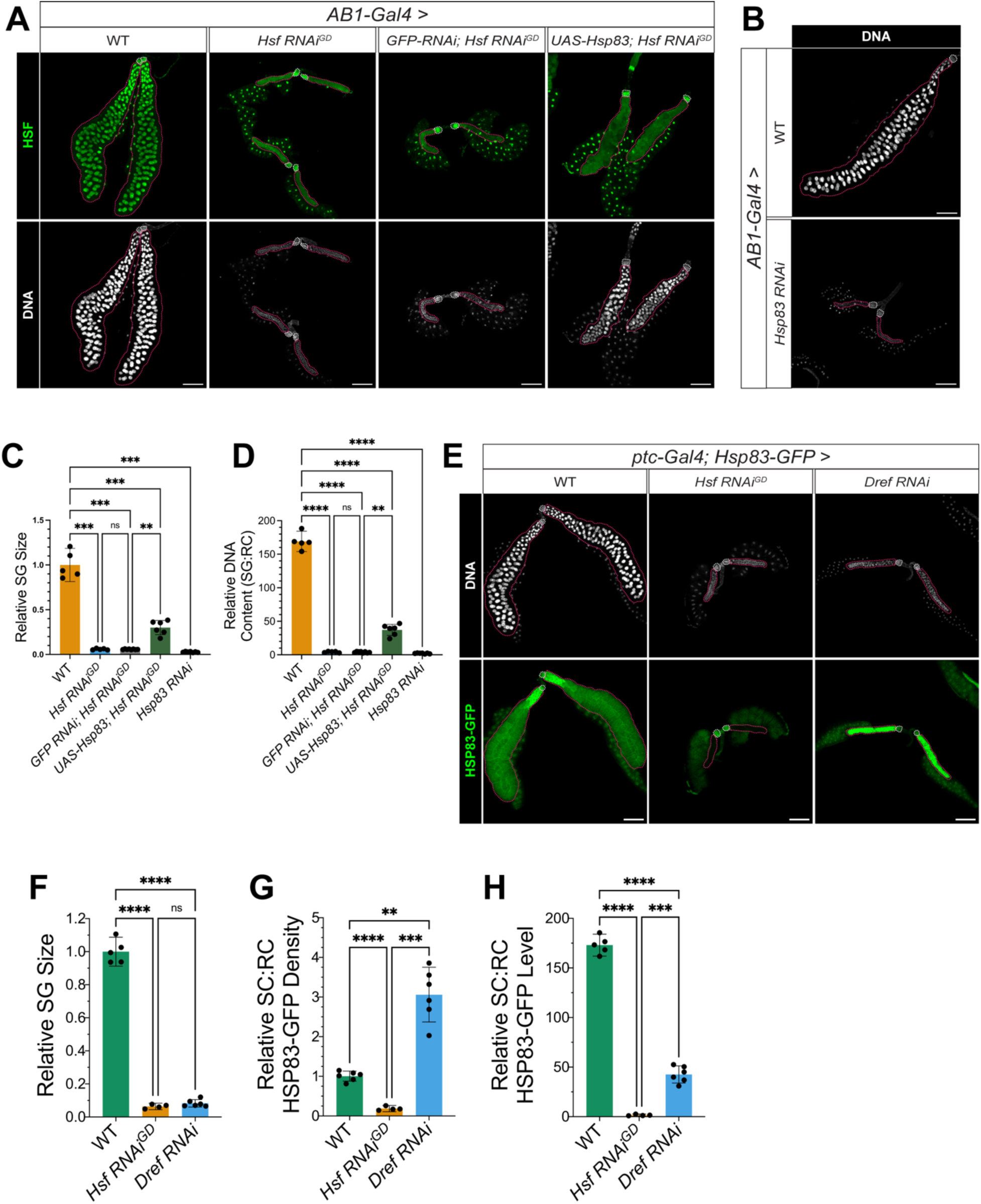
*Hsf* is necessary for salivary gland growth by maintaining basal HSP83 expression. (A) Immunostaining for HSF (green) and Hoechst staining for DNA (white) in the salivary gland (SG) of wandering 3^rd^ instar larvae at ∼120 h after egg laying (hAEL) in control and salivary gland-specific knockdown of *Hsf* (*Hsf RNAi*) animals, with or without HSP83 re-expression (*UAS-Hsp83*). The SGs and imaginal ring cells are outlined with red and white dashed lines, respectively. Scale bar, 150 μm. (B) Hoechst staining for DNA (white) in the salivary gland (SG) of wandering 3^rd^ instar larvae at ∼120 h after egg laying (hAEL) in control and *Hsp83 RNAi* animals. The SGs and imaginal ring cells are outlined with red and white dashed lines, respectively. Scale bar, 150 μm. (C) Quantitation of relative SG size, in control, *Hsf RNAi*, with or without HSP83 re-expression, and *Hsp83 RNAi* SGs from (A) and (B). n = 5-7 SGs per genotype. (D) Quantitation of DNA content per salivary gland, normalized to imaginal ring cells, in control, *Hsf RNAi*, with or without HSP83 re-expression, and *Hsp83 RNAi* SGs from (A) and (B). n = 5-7 SGs per genotype. (E) Hoechst staining for DNA (white) and HSP83–GFP (green) in the salivary glands of wandering 3^rd^ instar larvae at ∼120 h after egg laying (AEL) in control, *Hsf RNAi^GD^*, and *Dref RNAi* animals. The SGs and imaginal ring cells are outlined with red and white dashed lines, respectively. Scale bar, 150 μm. (F) Quantitation of relative SG size in control, *Hsf RNAi^GD^*, and *Dref RNAi* animals from (E). n = 4-6 SGs per genotype. (G) Quantitation of relative HSP83–GFP density (total SG signal normalized to SG area and then to the imaginal ring cells) from (E). n = 4-6 SGs per genotype. (H) Quantitation of relative HSP83–GFP level (total SG signal normalized to the imaginal ring cells) from (E). n = 4-6 SGs per genotype. Statistical significance was determined using Brown-Forsythe and Welch ANOVA followed by Dunnett’s multiple comparisons test. Data are presented as mean ± 95% CI. P < 0.05 (*), P < 0.01 (**), P < 0.001 (***), P < 0.0001 (****).

### HSF Regulates Wing Disc Development and Hemocyte Function via HSP83

Beyond endoreplication-driven growth, tissues also expand via proliferation (Hariharan, 2015). We therefore asked whether HSF is required in a proliferative tissue, the larval wing disc. During larval stages, the wing disc undergoes approximately 10 rounds of cell division, resulting in a 1,000-fold increase in cell number (Tripathi and Irvine, 2022). HSP83 is expressed in larval wing discs and has been implicated in cell-cycle exit (Bageritz et al., 2019; Bandura et al., 2013). Knocking down *Hsf* in wing discs using the *MS1096-Gal4* driver resulted in blistered, smaller, and shriveled adult wings, defects that were rescued by restoring HSP83 expression (Fig. 7A, B). These findings indicate that *Hsf* is required for wing development, in part through HSP83.

**Fig. 7.**
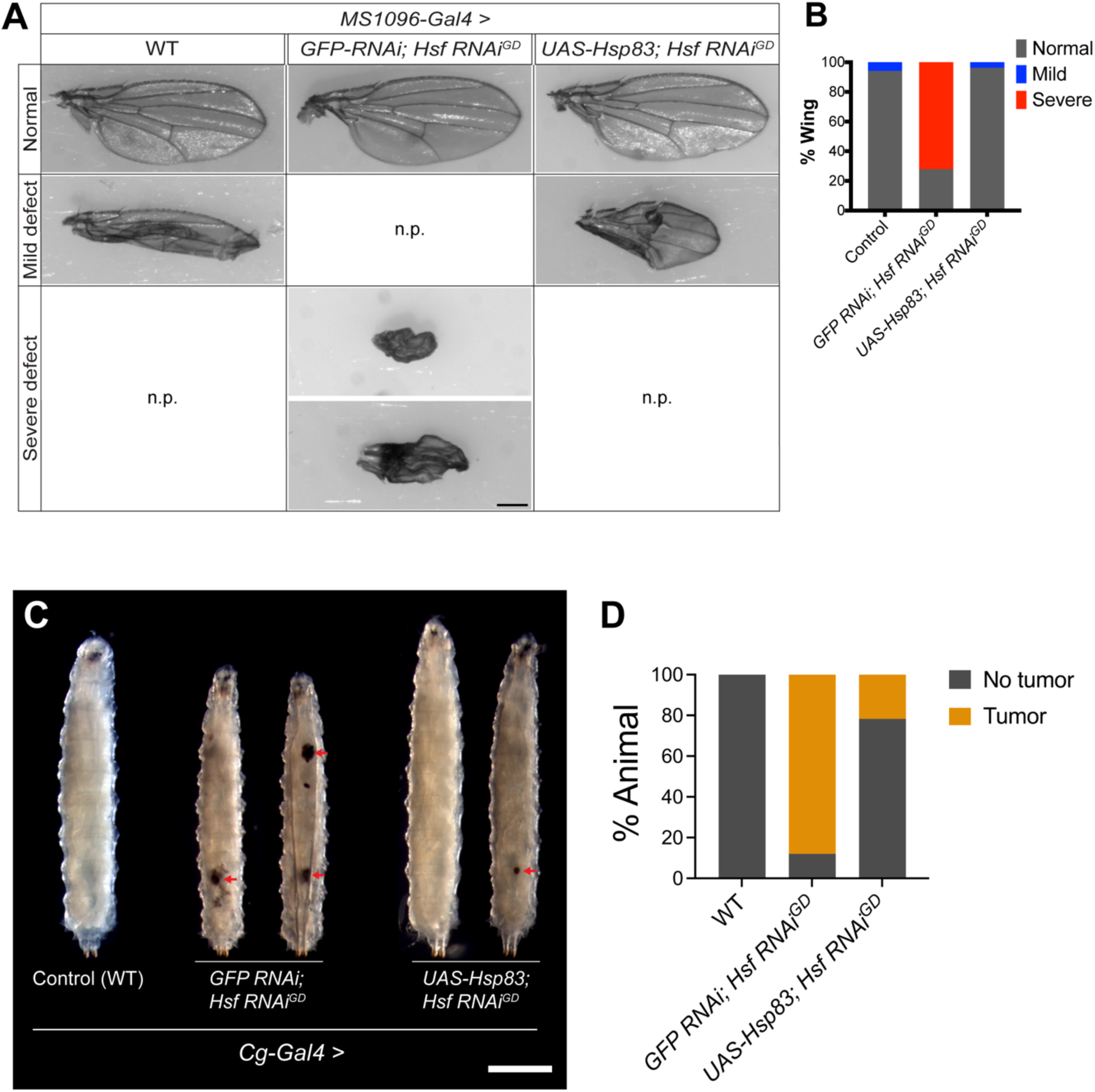
*Hsf* is necessary for normal wing Disc development and hemocyte functions via HSP83. (A) Representative images of adult wings from control and wing-disc specific knockdown of *Hsf* (*Hsf RNAi)* animals, with or without HSP83 re-expression (*UAS-Hsp83*). Wing phenotypes are categorized in three groups based on severity: Normal (no defects), mild defects, and severe defects. n.p., no such phenotype in that genotype. Scale bar, 0.5 mm. (B) Quantitation of wing phenotype distribution in control and *Hsf RNAi* animals, with or without *UAS-Hsp83* from (A). Color coding: normal (grey), mild defects (blue), and severe defects (red). (C) Representative images of control and hemocyte/fat body-specific knockdown of *Hsf* (*Hsf RNAi)* larvae, with or without HSP83 re-expression (*UAS-Hsp83*). Melanotic tumors are indicated by red arrows. Scale bar, 1 mm. (D) Quantitation of larvae with or without melanotic tumor in control and *Hsf RNAi* animals, with or without *UAS-Hsp83* from (A). Color coding: no tumor (grey) and tumor (yellow).

We also examined the role of *Hsf* in larval hemocytes, blood cells involved in immune surveillance and defense against pathogens and parasites (Meister and Lagueux, 2003; Williams, 2007). During embryonic and larval development, hemocyte numbers primarily increase through proliferation (Jung et al., 2005; Makhijani et al., 2011), and dysregulated proliferation can result in melanotic tumors (Minakhina and Steward, 2006; Ponrathnam et al., 2021). HSP83 is also expressed in hemocytes (Tattikota et al., 2020). To test if *Hsf* is necessary for hemocyte development, we knocked down *Hsf* in hemocytes using the strong fat-body/hemocyte driver *Cg-Gal4* (Ponrathnam et al., 2021). *Hsf* knockdown caused melanotic tumors in 88% of larvae, while restoring HSP83 expression partially rescued the phenotype, reducing the incidence to 33% (Fig. 7C, D). These results demonstrate that *Hsf* is essential for proper hemocyte development or proliferation via HSP83. Together, we conclude *Hsf* is also required for the proliferative tissue during larval development and acts at least in part through HSP83.

## MATERIALS AND METHODS

### Reagents and resources

See Table S1 for detailed information on the flies and reagents used in the studies and Table S2 for the specific genotypes used in each figure/panel.

### Fly stocks

*Oregon R* was used as the wild-type strain. *Hsf^1^*(Bloomington *Drosophila* Stock Center [BDSC], #5491) and *Hsf^03091^*(BDSC #11271) are *Hsf* loss-of-function alleles. *Df(2R)ED3610* (BDSC #9066) is a deficiency that removes *Hsf* as well as other neighboring genes. *Hsf-Halo,* a gift from Dr. Carl Wu, was used as *Hsf* rescue construct (Tang et al., 2022). *Hsf-RNAi^GD^* (Vienna *Drosophila* Resource Center [VDRC], #GD-37699) and *Hsf-RNAi^KK^* (VDRC #KK-108851) were used for RNAi-mediated *Hsf* knockdown. *phm-Gal4* (BDSC #80577) was used as a prothoracic gland driver. *UAS-CD8-RFP*, a gift from Dr. Dan Bergstralh, was used to express membrane-targeted RFP to label prothoracic gland membranes. *Hsp83-GFP*, a gift from Dr. Valeria Palumbo, was used to assess HSP83 protein levels in the prothoracic gland, salivary gland, and whole larvae (Tariq et al., 2009; Palumbo et al., 2020). *Dref-RNAi* (BDSC #31941), *Hsp83-RNAi* (VDRC #GD-7716), and *GFP-RNAi* (BDSC #9331) were additional shRNA lines used in this study; *GFP-RNAi* served as a control in RNA inference experiments (see below). *UAS-Hsp83* (BDSC #58469) was used to re-express HSP83 under *Hsf* knockdown conditions. *AB1-Gal4* (BDSC #1824) and *ptc-Gal4* (BDSC #2017) were used as salivary gland drivers. *Cg-Gal4* (BDSC #7011) was used as a hemocyte driver (also active in fat body and lymph gland). *MS1096-Gal4* (BDSC #8860) was used as a dorsal wing disc driver. *traffic-jam-Gal4*, a gift from Dr. Dan Bergstralh, was used as a follicle cell driver. *koi^84^/CyO,GFP*, a gift from Dr. Janice A. Fischer, was used as a source of a green fluorescent second chromosome balancer (Kracklauer et al., 2007). *UAS-TARDBP.J/CyO,P{2xTb1-RFP}CyO* (BDSC #51371) was used as the source of a red fluorescent second chromosome balancer. We used FlyBase to find information on phenotypes/function/stocks/gene expression (Öztürk-Çolak et al., 2024).

Flies used in all experiments were maintained on molasses/agar/yeast/malt extract/corn flour/soy flour food at room temperature except as noted below. For larval developmental stage experiments (Fig. 1, Fig. S1, and Fig. S3), larvae were raised on freshly prepared food consisting of 2.5 g dry yeast, 2.5 g Formula 4-24 Instant *Drosophila* Medium (Carolina Biological Supply #173200, and gift from Dr. John Jaenike), and 15 mL deionized water. For prothoracic gland and salivary gland experiments, flies were raised on food adapted from the Okamoto food (Okamoto et al., 2018): 30 g of agar gelidium (Mooragar #41084), 250g brown brewer’s dry yeast, 500 g Clintose dextrose (Fisher #NC178886), 350 g Quaker enriched cornmeal (Walmart), 13 g of methyl paraben (Methyl-p-Hydroxybenzoate; VWR #IC10234105), 30 mL propionic acid (Fisher #AC149300010), 50 mL ethanol, and 5 L deionized water.

### Developmental staging

*Hsf* loss-of-function mutants and deficiency lines were maintained over the green-fluorescent second-chromosome balancer CyO,GFP, which was visualized in larvae using a NIGHTSEA Stereo Microscope Fluorescence Adapter. Crosses using at least 40 males and females were set up in small cages with 6 cm apple juice plates to collect eggs for 2 h of the desired *Hsf* genotypes. To synchronize larval development, plates were gently washed with deionized water 24 h after egg laying to remove larvae that had already hatched. One hour later, at least 30 newly hatched larvae of the desired genotype were selected using NIGHTSEA fluorescence and transferred to freshly prepared food paste (yeast/instant fly food/water) on 6 cm apple juice plates designated as 0 h after egg hatching, AEH). Plates were maintained in a humidified chamber at 25 °C. Every 6 to 24 hours after the collection, larvae were washed off using 3M NaCl to assess developmental stage based on spiracle and tracheal tube morphology. After staging, larvae were transferred to a fresh food plate. Each experiment was repeated three times on three different days.

For percent pupariation assays, at least 30 larvae were synchronized as described above and transferred to tubes containing adapted Okamoto food. The number of animals that had entered the pupal stage (white or brown pupae) was counted every 4 h from 116 to 148 h after egg laying (AEL). Percent pupariation was calculated as the number of pupae divided by the total number of larvae initially transferred to each tube. Each experiment was repeated at least three times on three independent days.

### RNA Interference

RNA interference (RNAi) experiments were performed by crossing approximately 15 females from the appropriate Gal4 driver line with 15 males from the corresponding UAS-RNAi line. For salivary gland and prothoracic gland knockdown experiments, crosses were maintained at 25 °C until dissection. For wing-disc knockdown experiments, crosses were maintained at 29 °C until eclosion of the F1 progeny, at which point wing phenotypes were assessed. For follicle cell knockdown experiments, crosses were maintained at 25 °C until eclosion of the F1 progeny. Newly eclosed adults were then shifted to 29 °C for at least five days prior to ovary dissection. For HSP83 re-expressing experiments, UAS-*GFP-RNAi* was used in the control group lacking UAS-*Hsp83* to keep the total number of UAS targets consistent between the control and experimental groups.

### Fixation and immunostaining

Flies used for prothoracic gland and salivary gland experiments were synchronized as described above and raised on adapted Okamoto food until the wandering 3^rd^ instar larval stage, at which point dissections were performed. For *Hsp83* knockdown in the prothoracic gland, larvae died during the 2^nd^ instar stage; therefore, dissections were performed using 2^nd^ instar larvae. Prothoracic glands and salivary glands were dissected from larvae in phosphate-buffered saline (PBS) and fixed in 4% paraformaldehyde prepared in PBS-T (PBS containing 0.1% Triton X-100) for 15 min at room temperature (RT). Ovaries were fixed in 6% formaldehyde in 100 mM KH_2_PO_4_/K_2_HPO_4_ pH 6.8, 450 mM KCl, 150 mM NaCl, 20 mM MgCl_2_, with vigorous shaking for 20 minutes at RT (Wang and Lindquist, 1998). Tissues were washed three times in PBS-T for 15 min each at RT and then blocked overnight at 4 °C in blocking buffer (10% bovine serum albumin, 0.1% Triton X-100, and 0.02% sodium azide in PBS). Tissues were subsequently incubated with primary antibodies diluted in blocking buffer overnight at 4 °C. After primary antibody incubation, tissues were washed three times in PBS-T for 15 min each at RT and then incubated with secondary antibodies diluted in blocking buffer overnight at 4 °C. Tissues were washed three times in PBS-T for 15 min each at RT, followed by DNA staining with Hoechst 33342 (Thermo Fisher Scientific) at a 1:1000 dilution in blocking buffer for 30 min at RT. After three additional washes in PBS-T for 15 min each at RT, tissues were mounted in Aqua-Poly/Mount (PolySciences).

The following primary antibodies were used at the indicated dilutions: rabbit anti-HSF (1:1000; gift from Dr. Susan Lindquist, originally from Dr. Tim Westwood) (Westwood et al., 1991), rabbit anti-Hsp70 (7FB; 1:1000; gift from Dr. Susan Lindquist; specifically recognizes the heat-inducible Hsp70 family member in *Drosophila*) (Velazquez et al., 1983), and rabbit anti-Dib (1:1000; gift from Dr. Naoki Yamanaka). Goat anti-rabbit IgG Alexa Fluor 488 and Alexa Fluor 633 (Invitrogen) were used as secondary antibodies (1:10000) to detect primary antibodies. Secondary antibodies were pre-cleared by overnight incubation at 4 °C with wild-type larval tissues to reduce non-specific staining.

### Western Blot

Newly hatched (0 h AEH) and 24 h AEH wild-type and *Hsf* null larvae were heat-fixed in 1x Triton Salt Solution for 1 minutes and boiled in Laemmli buffer (Bio-RAD, Hercules, CA) for 15 minutes. Proteins were separated using 4-15% SDS-PAGE gels (Bio-RAD) and transferred to PVDF membranes (Immobilon-FL, EMD Millipore, Burlington, MA). Transfers were performed in Towbin buffer (10% 10x Tris-Glycine solution, 20% Methanol) at 80V for 30 mins. Immunodetection was done using the following primary antibodies – rabbit anti-HSP90 (1:1000) (Cell Signaling, Danvers, MA) and mouse anti-α-Tubulin (1:1000) (Cell Signaling, Danvers, MA) and secondary antibodies – IRDye® 800CW Goat anti-Rabbit IgG and IRDye® 680RD Goat anti-Mouse IgG (1:10,000) (Li-COR, Lincoln, NE). The same membranes were probed with the indicated primary antibody and mouse anti-α-tubulin as a loading control. Images were acquired using Li-COR Odyssey CLx, and fluorescence was quantified using Image StudioLite Software.

### Microscopy

Prothoracic gland and salivary gland images were acquired using an Andor Dragonfly spinning-disk confocal microscope at the University of Rochester High Content Imaging Core with 63× (NA = 1.49) and 10× (NA = 0.45) objectives, respectively. Female ovaries were imaged using a Leica SP5 confocal microscope with a 20× objective (NA = 0.75). Mouth hook imaging (Fig.1) and epifluorescence imaging of larvae (Fig. 2) were performed on a Nikon Eclipse E600 microscope using 20× (NA = 0.50) and 4× (NA = 0.10) objectives, Spot Insight imaging software, and a Diagnostics Instruments 14.2-megapixel color mosaic camera. All other larval and wing disc images were acquired using a Moticam 2000 camera. All images were processed using FIJI (NIH) and assembled in Adobe Illustrator.

### DNA Quantification

For the prothoracic gland, three-dimensional reconstruction of z-stack images was performed to determine the total number of nuclei. Total DNA content was calculated by summing signal intensity across all slices of the DAPI channel after background subtraction. DNA content per nucleus was calculated by dividing the total DNA content by the number of nuclei and then normalized to that of the corpus allatum, a neighboring endocrine gland not targeted by the *phm-Gal4* driver. For the salivary gland, DNA content per gland was calculated by dividing the total DNA content of the gland by the total DNA content of the imaginal ring cells, a small population of diploid precursor cells located at the proximal end of the salivary gland that are not targeted by the *AB1-Gal4* or *ptc-Gal4* drivers.

### Gland size quantification

Prothoracic gland size was determined by calculating the area of the maximum-intensity projection of the CD8-RFP signal. Salivary gland size was determined based on the tissue outline defined by background fluorescence in the HSF immunostaining. Relative gland size was calculated by normalizing to the size of wild-type glands.

### HSP83 level quantification

For the prothoracic gland, salivary gland, and whole larvae, HSP83 levels were quantified using a *Hsp83-GFP* reporter line (Tariq et al., 2009; Palumbo et al., 2020). GFP level was defined as the total GFP fluorescence intensity measured across all z-slices in the GFP channel after background subtraction within the gland or larval region. GFP density was calculated by dividing the GFP level by the area of the gland or larva. For the prothoracic gland, both GFP level and GFP density were normalized to those of the corpus allatum. For the salivary gland, both parameters were normalized to those of the imaginal ring cells.

### 20E feeding

Larvae with *Hsf* knockdown in the prothoracic gland were raised on standard food (dry yeast and instant fly food) until 48 h after hatching, after which they were transferred to either standard food or food supplemented with 20E. The number of pupae was subsequently recorded. Larvae with *Hsp83* knockdown in the prothoracic gland were raised on standard food or food supplemented with 20E from hatching, and developmental stage was scored every 24 h as described above. Food supplemented with 20E was made by mixing 100 mg dry yeast, 100 mg instant fly food, 380 μL deionized water, and 10 mM 20E in ethanol.

### Larval Foraging Assay

Larvae with *Hsp83* knockdown in the prothoracic gland were raised on standard food (dry yeast and instant fly food) or food supplemented with 20E made as described above until 36 h after hatching. Larvae were removed from the food using 3 M NaCl. For the foraging assay, a blob of yeast paste was placed at the center of a 6-cm apple juice plate. Larvae were gently transferred to the edge of the plate, and the number of larvae remaining outside the food source was recorded every 5 min.

### Heat shock assay

Heat shock experiments were performed by placing 30 wandering 3^rd^ instar larvae into a scintillation vial and submerging the vial in a 42 °C water bath for 20 min. Following heat shock, the vial was immediately placed on ice to rapidly cool the larvae. Larvae were then immediately dissected in PBS and processed for fixation and immunostaining using the protocols described above.

### Statistics

All statistical analyses were performed using GraphPad Prism 10 (v10.6.1). Details of statistical tests and sample sizes are provided in the figure legends.

## DISCUSSION

### HSF regulates *Drosophila* developmental growth via HSP83

Here we show that Heat Shock Factor (HSF) supports developmental growth in multiple *Drosophila melanogaster* tissues. *Hsf* null larvae die at early 2^nd^ instar, and tissue-restricted *Hsf* depletion reveals organ-specific requirements: reduced endoreplication and growth in the prothoracic and salivary glands and in follicle cells; adult wing defects after larval imaginal-disc knockdown; and melanotic masses upon hemocyte knockdown. These observations align with the previous finding that *Hsf^1^* germline clones arrest during oogenesis at stages 5-6, exhibiting endoreplication and growth defects (Jedlicka et al., 1997).

We also provide evidence that in these tissues HSF maintains the basal expression of HSP83, based on either directly measuring HSP83 levels and/or rescuing the growth defects by overexpressing HSP83. Our results agree with prior evidence that HSF sustains basal HSP83: an RNA aptamer targeting HSF reduces *Hsp83* mRNA by ∼50% in the salivary gland (Salamanca et al., 2011), and *Drosophila* Kc cells ChIP maps strong HSF enrichment upstream of the *Hsp83* promoter (Gonsalves et al., 2011). At first glance, this finding seems at odds with Jedlicka et al. 1997, who argued that HSF’s developmental functions during oogenesis and early larval stages are not mediated by HSP induction. Several differences may explain these discrepancies. First, Jedlicka et al. analyzed the temperature-sensitive *Hsf^4^* allele at the non-permissive temperature of 29 °C rather than *Hsf* nulls; because *Hsf^4^* mutants lack a functional heat shock response, lethality at 29 °C might in part reflect chronic heat stress and proteotoxicity rather than a basal developmental role. Second, during oogenesis, they assessed HSP83 with an HSP83–lacZ reporter and X-Gal staining in *Hsf* germline clones; intact HSP83 expression in surrounding follicle cells (unaffected in the clones) would be expected to dilute or mask potential signal loss in the nurse cells at light-microscopy resolution. Third, quantitative measurements were not reported; the slightly weaker central X-Gal staining in *Hsf* follicles is compatible with a modest decrease. Finally, it is possible that HSP83’s dependence on HSF may be tissue- and stage-specific.

### HSF is critical for proteostasis during animal development

How animals achieve large, rapid increases in organ and tissue size over short developmental windows is a central question in developmental biology. Developing tissues have to sustain adequate protein-folding and maturation capacity during growth-driven protein synthesis. Developmental growth couples increased translation with enhanced ribosome biogenesis, imposing heavier loads on the folding machinery (Brown and Dawid, 1968; Surrey et al., 1979; Falcon et al., 2022). In secretory tissue, cells cope by expanding endoplasmic reticulum folding capacity via the unfolded protein response during growth and differentiation (Cornejo et al., 2013; Hetz and Papa, 2018; Reimold et al., 2001). By contrast, how developing tissues scale nucleo-cytoplasmic proteostasis with surging translation and their regulators remains incompletely understood.

Our results suggest that HSF is a candidate regulator of nucleo-cytoplasmic proteostasis during animal development by maintaining the basal expression of the molecular chaperone HSP83/HSP90, known to maintain proteostasis. This requirement for HSF in developmental growth appears conserved across eukaryotes. In *Caenorhabditis elegans*, *Hsf-1* null larvae arrest and at the early larval stage, and germline-specific HSF depletion impairs mitotic proliferation of germline progenitors (Li et al., 2016; Edwards et al., 2021). In mice, *Hsf1* deficiency causes postnatal growth retardation (Xiao et al., 1999). A principal function underlying this requirement is maintenance of proteostasis. In *C. elegans* larvae, 14 of the 22 essential HSF-regulated genes are involved in protein folding, including heat shock proteins (HSP90, HSP110, HSC70) and components of the chaperonin-containing T-complex (Li et al., 2016). In the germ line, HSF depletion downregulates 9 constitutively expressed chaperones, including HSP-90 and HSC-70 (Edwards et al., 2021). Maintaining basal expression of HSPs appear to be an ancestral function as in *S. cerevisiae*, *Hsf1* sustains basal expression of 18 chaperone genes, and re-expression of just two—*Hsp70* and *Hsp90*—was sufficient to restore viability upon *Hsf1* depletion (Solís et al., 2016).

Unlike fungi and invertebrates, which encode a single *Hsp90*, mammals have two paralogs—*Hsp90α* and *Hsp90β*—encoded by separate genes (Csermely et al., 1998; Prodromou, 2016; Subbarao Sreedhar et al., 2004). Both are expressed basally *in vivo*, with *Hsp90α* often stress-inducible and *Hsp90β* more constitutive. The extent to which their basal levels depend on HSF during development appears context dependent. For example, in mouse embryos at embryonic day 11.5–13.5, *Hsf1* knockout did not measurably alter *Hsp90α* protein levels (Xiao et al., 1999). By contrast, oocytes from *Hsf1* null females display defective meiosis accompanied by significantly reduced HSP90α, and pharmacological inhibition of HSP90α produces similar phenotypes (Metchat et al., 2009). For HSP90β, available data are limited: one report states that HSP90β is unchanged in *Hsf1* null embryos, yet the single image provided appears reduced and no quantification was reported (Xiao et al., 1999).

Together, cross-species evidence supports a conserved, tissue- and stage-dependent role for HSF in sustaining proteostasis during development. This notion is consistent with our finding in *Drosophila* that HSF maintains basal HSP83 (HSP90) to support growth. Although we did not assess other HSPs in this study, the robust rescue of HSF-depletion phenotypes by HSP83 re-expression suggests that HSP83 is a principal route by which HSF promotes developmental proteostasis, without excluding contributions from additional chaperones. Consistent with this view, in *Drosophila*, HSP83 has been implicated in regulating proteasomal activity (Choutka et al., 2017), neuronal cell reactivation (Huang and Wang, 2018), microtubule polymerization (Palumbo et al., 2020), spermatogenesis (Yue et al., 1999), ecdysone receptor activation (Arbeitman and Hogness, 2000), and nutrient sensing during developmental (Ohhara et al., 2021).

### Regulation of HSF activity during development

The requirement for HSF in maintaining basal HSP expression raises key questions about how HSF is regulated in vivo during development. Upon stress, it is theorized that HSF is phosphorylated, trimerizes, translocates to the nucleus, and binds to conserved DNA sequence motifs known as the heat shock elements (HSEs) in the promoters of chaperone genes, thereby initiating their transcription. Whether these regulatory steps also happen for HSF to support developmental growth is a remaining question.

Is trimerization required for basal activity? Trimerization is essential for stress-induced HSP upregulation (Baler et al., 1993; Sarge et al., 1993), but appears dispensable for yeast growth under non-stress conditions: HSF1 lacking the trimerization domain supports normal growth at 28 °C yet fails under heat shock at 38 °C (Hashikawa et al., 2007). One possibility is that monomeric HSF can sustain basal Hsp90 transcription by binding directly to the heat-shock elements (HSEs) at the promoter. Given the very low in-vitro DNA-binding affinity of monomeric HSF, additional cofactors or chromatin context may facilitate binding in vivo (Jaeger et al., 2014).

What activates HSF under non-stress conditions? Traditionally, HSF1 is thought to remain inactive in the cytoplasm and only translocate to the nucleus upon stress-induced trimerization (Baler et al., 1993; Sarge et al., 1993). However, more recent studies using immunostaining and fluorescent protein tagging indicate that HSF is predominantly nuclear in *Drosophila*, *C. elegans*, rodents, and humans (Fairfield et al., 2002; Joutsen et al., 2024; Mercier et al., 1999; Morton and Lamitina, 2013; Wang and Lindquist, 1998; Yao et al., 2008). This constitutive nuclear localization suggests that HSF activity is regulated by additional mechanisms beyond nuclear translocation, allowing context-specific activation under non-stressed conditions. Distinct pools of HSF may exist, with a constitutively nuclear, DNA-bound fraction providing basal transcriptional activity.

Do cofactors specify developmental outputs? Tissue-specific Hsp90 dependence on HSF in mammals may reflect cofactor modulation of HSF activity. For example, in HuH7 hepatoma cells, interferon-” treatment induces the activation of STAT-1 which physically interacts with HSF to synergistically increase Hsp90β expression (Stephanou et al., 1999). Similarly, IL-6 promotes Hsp90β expression through STAT-3 activation; however, in this context, STAT-3 antagonizes HSF activity (Stephanou et al., 1998). These observations may be explained by the overlapping binding sites for STATs and HSF in the Hsp90β promoter (Stephanou and Latchman, 1999). In addition, the basal expression of Hsp90 can be co-regulated by other transcription factors, including Sp1 family transcription factors (Fu et al., 2023). In *C. elegans*, developmental HSF-1 activation requires E2F/DP binding to a GC-rich motif that facilitates HSE recognition (Li et al., 2016). Together, these observations point to a cofactor-defined, context-dependent HSF program that could explain tissue- and stage-specific requirements during development.

### HSF and Diseases

The importance of *Hsf* in regulating proteostasis under non-stress conditions suggests a broader role in protein folding diseases and aging. In mammalian cancer, HSF promotes the growth of many tumors by driving a transcriptional program that is distinct from that following stress but HSP90 and certain other heat shock proteins (Dai et al., 2007; Mendillo et al., 2012; Santagata et al., 2013). In Parkinson’s disease models, pharmacological activation of HSF reduces α-synuclein aggregation by inducing HSP70 expression (Ekimova et al., 2018; Liangliang et al., 2010). Similarly, in polyglutamine (PolyQ) disease models, HSF levels negatively correlate with pathogenic aggregate formation, and overexpression of constitutively active HSF suppresses aggregation by upregulating major heat shock protein genes (Fujimoto et al., 2005; Kondo et al., 2013). HSF is also linked to aging, as the progressive decline of proteostasis and the heat shock response is a hallmark of aging (López-Otín et al., 2023). Notably, overexpression of HSF extends lifespan in *Caenorhabditis elegans* and *Drosophila melanogaster* (Glastad et al., 2023; Morley and Morimoto, 2004).

In addition to its role in maintaining proteostasis, HSF may also exert non-transcriptional functions under non-stress disease conditions. In mammalian tumors, HSF1 is inactivated by AMP-activated protein kinase (AMPK) through phosphorylation, after which it physically interacts with AMPK to suppress its activity; in mice, this interaction promotes a lipogenic phenotype and tumor growth independently of HSF1’s canonical transcriptional activity (Su et al., 2019). Moreover, in HeLa cells, HSF1 enhances c-MYC–mediated transcription by forming a complex with c-MYC and MAX, independent of its own DNA-binding activity (Xu et al., 2023). These findings suggest that HSF can influence cellular physiology through mechanisms beyond direct transcriptional regulation of heat shock genes. Together, these studies suggest that functions of HSF under non-stress condition are diverse and context-dependent, from transcriptional regulation of proteostasis to non-transcriptional regulation of key signaling and transcriptional networks. Given these complexities, a full understanding of HSF1’s role in development may require the analysis of many distinct downstream targets. Our study shows that in many *Drosophila* tissues the regulation of HSP83 expression is a major critical target; reducing just basal HSP83 expression accounts for many, if not all, of the dramatic growth defects after loss of HSF. This critical role of HSP83 may therefore mask HSF’s modulation of other processes. Restoring HSP83 levels after HSF knockdown – like performed here – thus provides an opportunity to identify additional biologically important targets in the future.

### Limitations of the study

As HSF has many potential binding site in the genome, of which hundreds can be occupied at normal temperatures, it might promote growth and development by regulating hundreds of genes. Our data indicate that HSP83 is a major biologically relevant target as we can rescue multiple developmental defects upon HSF knockdown to a large extent by simply re-expressing HSP83. This is true for the prothoracic gland, salivary gland, follicle cells, the wing disc, and hemocytes. Not all of the knockdown phenotypes can be fully rescued by HSP83 re-expression. Future studies will have to determine whether partial rescue is the result of suboptimal HSP83 levels or reduced expression of other HSF targets. Whatever the case, our data identify an HSF – HSP83 axis that contributes to developmental growth.

## ACKNOWLEDGEMENTS

We are grateful to Dr. Susan Lindquist and Dr. Terry Orr-Weaver who inspired us to probe the role of HSF in development. We thank Jordan Aronowitz for initial HSF characterization in the Welte laboratory. Stocks obtained from the Bloomington *Drosophila* Stock Center (NIH P40OD018537) were used in this study. Additional transgenic fly stocks were obtained from the Vienna *Drosophila* Resource Center (VDRC, www.vdrc.at). We used FlyBase to find information on phenotypes/function/stocks/gene expression. We thank Dr. Susan Lindquist, Dr. Valeria Palumbo, Dr. Janice A. Fischer, Dr. Dan Bergstralh, Dr. Carl Wu, Dr. Naoki Yamanaka, and Dr. John Jaenkie for kindly sharing fly stocks and reagents. We thank Noah Reger for assistance with fly genotyping. We thank Brian Jencik for assistance with reagent ordering and administrative support. We thank the Welte Lab members for comments on the manuscript and discussions.

## AUTHOR CONTRIBUTIONS

J.J.T. and M.A.W. conceptualized the study and designed the experiments. J.J.T. performed all experiments except the wing disc and follicle cell experiments. A.S. and S.S. performed wing disc experiments. R.P.W. and E.F. performed follicle cell experiments. J.J.T. analyzed the data and wrote the original draft of the manuscript. J.J.T., A.S. and M.A.W. reviewed and edited the manuscript. M.A.W. acquired funding and supervised the study.

## DECLARATION OF GENERATIVE AI AND AI-ASSISTED TECHNOLOGIES

The first author wrote the first draft of this work and subsequently used ChatGPT to help polish the text for clarity. The authors then extensively reviewed and edited the manuscript and take full responsibility for the content of the publication.

