## Supplemental Materials for "*Drosophila* Heat Shock Factor (HSF) Regulates Developmental Growth by Maintaining the Basal Expression of HSP83/HSP90"

**Fig. S1.**

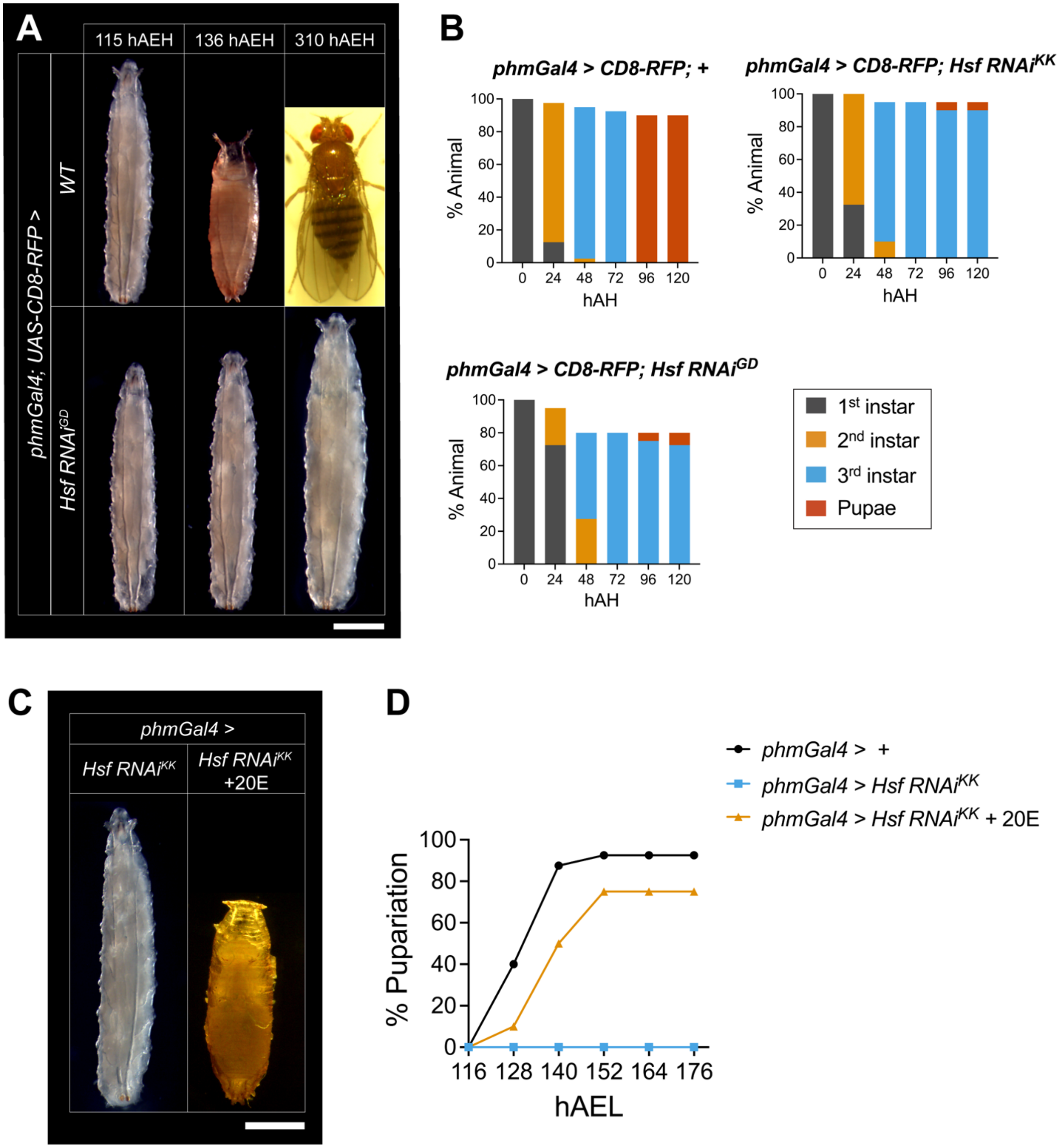
A
115 hAEH
136 hAEH
310 hAEH
phmGal4; UAS-CD8-RFP >
WT
Hsf RNAi^GD
B
phmGal4 > CD8-RFP; +
phmGal4 > CD8-RFP; Hsf RNAi^KK
phmGal4 > CD8-RFP; Hsf RNAi^GD
% Animal
hAH
0 24 48 72 96 120
0 20 40 60 80 100
1st instar
2nd instar
3rd instar
Pupae
C
phmGal4 >
Hsf RNAi^KK
Hsf RNAi^KK +20E
D
phmGal4 > +
phmGal4 > Hsf RNAi^KK
phmGal4 > Hsf RNAi^KK + 20E
% Pupariation
hAEL
116 128 140 152 164 176

**Fig. S1. Prothoracic gland (PG)-specific knockdown of *Hsf* causes a giant larva phenotype*.*** (A) Representative images of wild-type (WT) and PG-specific *Hsf* knockdown larvae (*Hsf RNAi*$^{GD}$) at 115, 136, and 310 h after egg hatching (hAEH). *Hsf* RNAi larvae remained arrested at the larval stage, whereas wild-type animals pupariated and eclosed. Scale bar, 1 mm. (B) Quantitation of developmental stage distribution and survival (%) in control and *Hsf* knockdown (*Hsf RNAi*$^{GD}$ and *Hsf RNAi*$^{KK}$) larvae. n = 40 larvae per genotype. (C) Representative images of *Hsf RNAi*$^{KK}$ larvae with or without 20E supplementation. Scale bar, 1 mm. (D) Percent pupariation over time in control and *Hsf RNAi*$^{KK}$ larvae with or without 20E supplementation from 116 to 148 h after egg laying (hAEL). n = 20-40 larvae per condition. 20E supplementation partially rescued the pupariation defect.

**Fig. S2.**

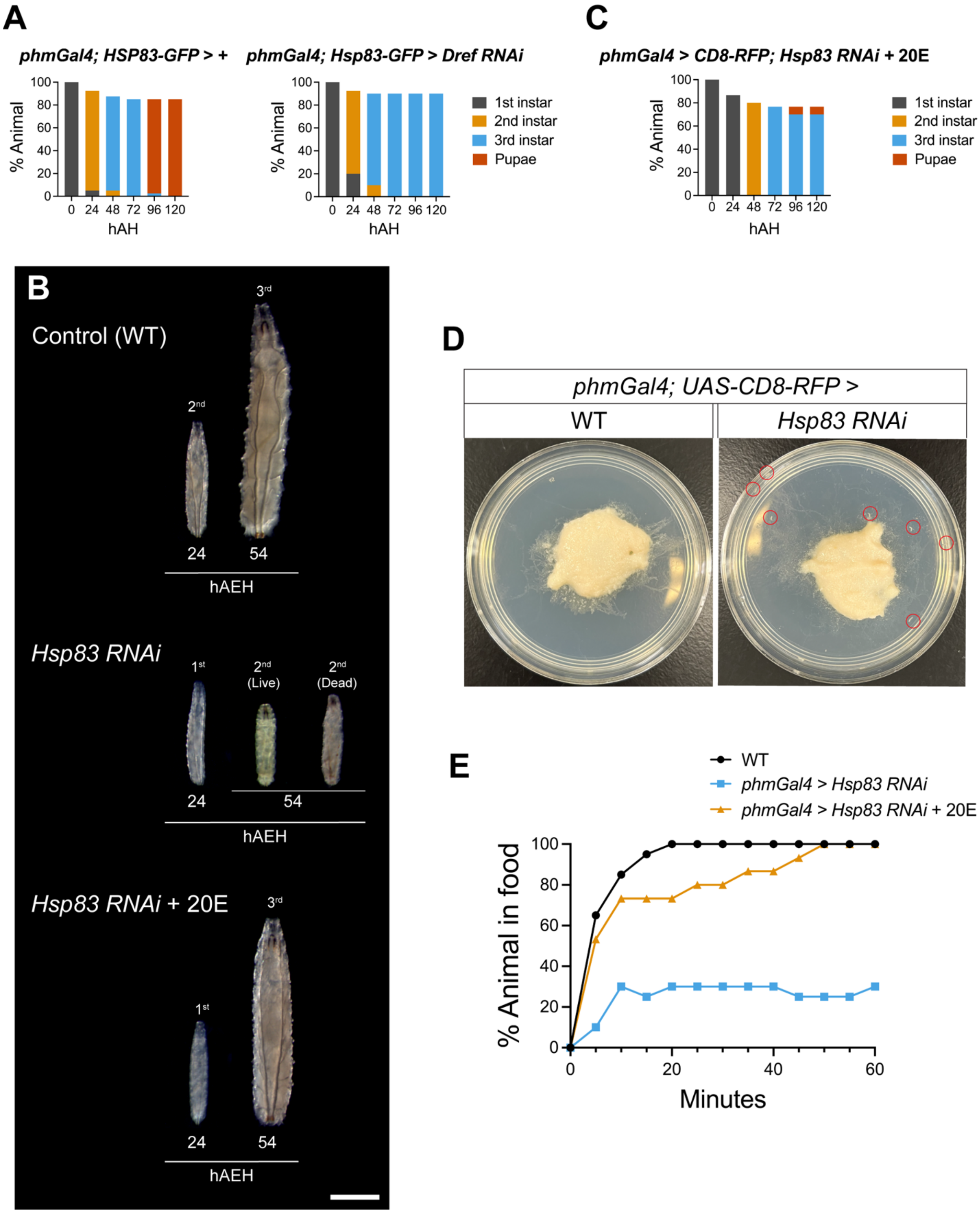
A
phmGal4; HSP83-GFP > +
phmGal4; Hsp83-GFP > Dref RNAi
% Animal
hAH
1st instar
2nd instar
3rd instar
Pupae
C
phmGal4 > CD8-RFP; Hsp83 RNAi + 20E
B
Control (WT)
hAEH
Hsp83 RNAi
(Live)
(Dead)
Hsp83 RNAi + 20E
D
phmGal4; UAS-CD8-RFP >
WT
Hsp83 RNAi
E
WT
phmGal4 > Hsp83 RNAi
phmGal4 > Hsp83 RNAi + 20E
% Animal in food
Minutes

**Fig. S2. Prothoracic gland (PG)-specific knockdown of *Hsp83* or *Dref* causes developmental arrest.** (A) Quantitation of developmental stage distribution and survival (%) in control and PG-specific *Dref* knockdown larvae. (B) Representative images of wild-type (WT) and PG-specific *Hsp83* knockdown larvae (*Hsf RNAi*$^{GD}$) at 24 and 54 h after egg hatching (hAEH). *Hsp83* RNAi larvae died in the 2$^{nd}$ instar stage. Scale bar, 1 mm. (C) Quantitation of developmental stage distribution and survival (%) in *Hsp83 RNAi* larvae with 20E supplementation (compare to Fig. 4E). 20E supplementation partially rescued the death during the 2$^{nd}$ instar stage, as indicated by the presence of 3$^{rd}$ instar larvae. n = 30 larvae. (D) Representative images of plates containing control (wild-type) and *Hsp83 RNAi* larvae. *Hsp83 RNAi* larvae, but not control larvae, wandered out of the food patch (circled in red). (E) One hour foraging assay showing the percentage of animals within the food patch over time in control and *Hsp83 RNAi* larvae with or without 20E supplementation. n = 15-20 larvae per condition.

1224 **Fig. S3.**

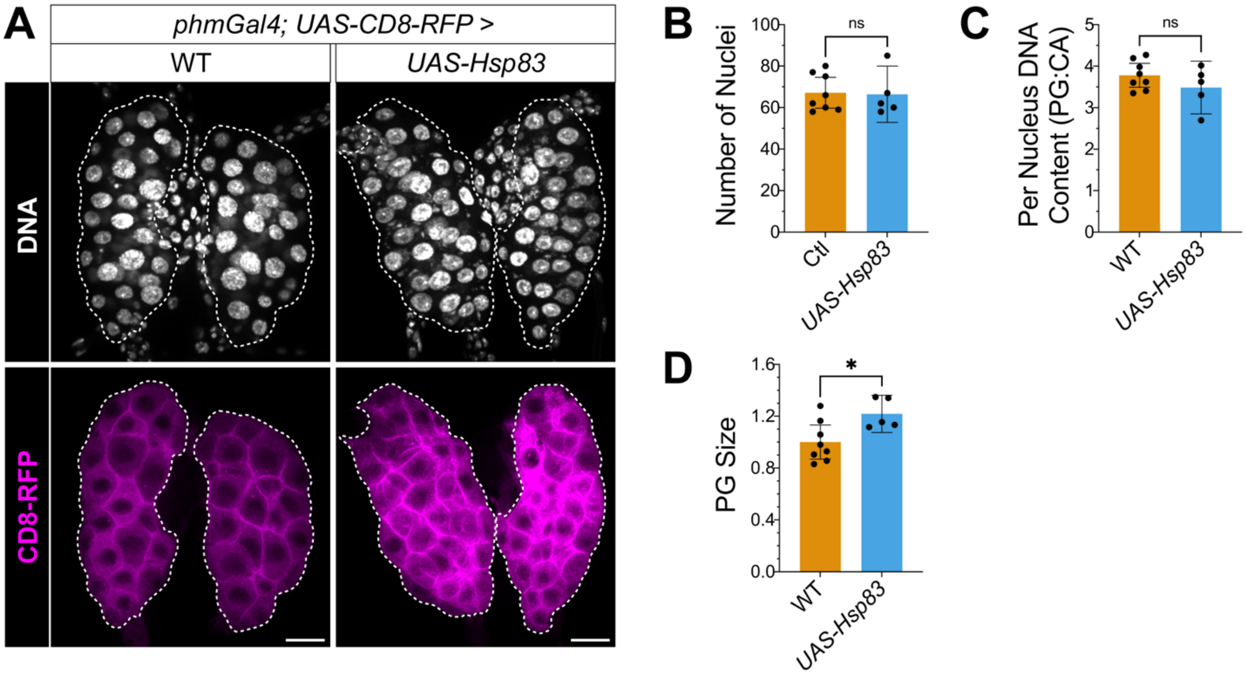


1225

1226

**Fig. S3. Overexpressing HSP83 on its own in the prothoracic gland (PG) minimally affects PG morphology.** (A) Hoechst staining for DNA (white) and CD8-RFP expression (magenta) in the prothoracic gland of wandering 3$^{rd}$ instar larvae at ~120 h after egg laying (hAEL) in control and HSP83 overexpression (*UAS-Hsp83)* animals. The prothoracic gland is outlined with white dashed lines. Scale bar, 20 μm. (B) Quantitation of the number of nuclei per PG in control and *UAS-Hsp83* PGs from (A). n = 8 and 5 PGs for each genotype. (C) Quantitation of DNA content per PG nucleus, normalized to corpus allatum (CA) nuclei in control and *UAS-Hsp83* PGs from (A). n = 8 and 5 PGs for each genotype. (D) Quantitation of relative PG size in control and *UAS-Hsp83* PGs from (A). n = 8 and 5 PGs for each genotype. Statistical significance was determined using Welch's t test. Data are presented as mean ± 95% CI. P < 0.05 (*), P < 0.01 (**), P < 0.001 (***), P < 0.0001 (****).

**Fig. S4.**

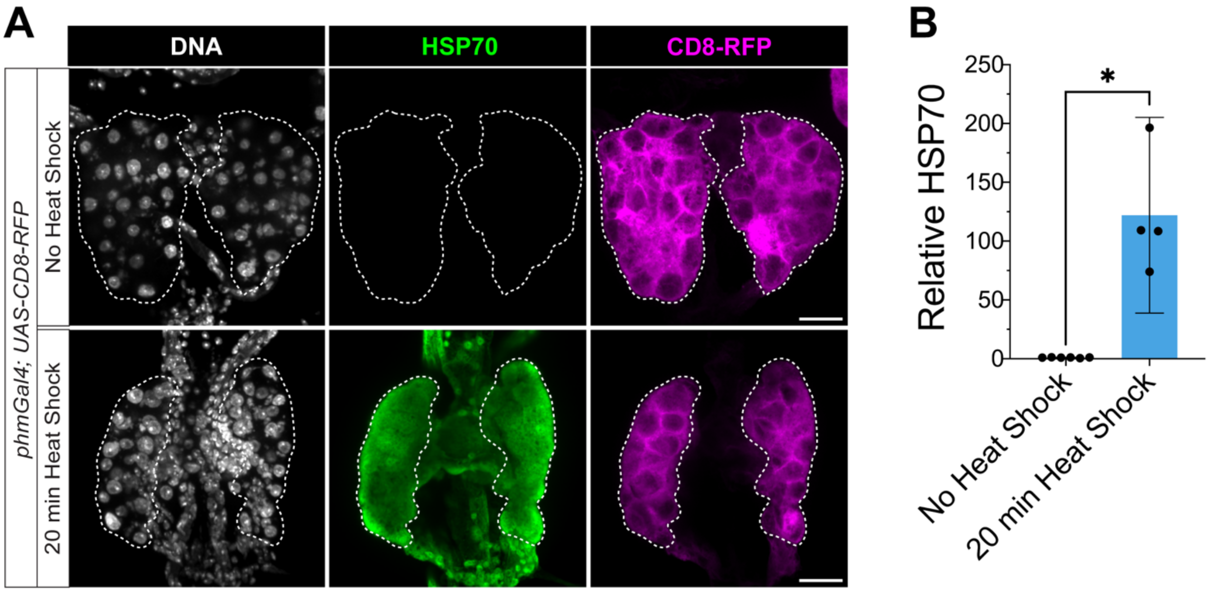
A
DNA
HSP70
CD8-RFP
phmGal4; UAS-CD8-RFP
No Heat Shock
20 min Heat Shock
B
Relative HSP70
250
200
150
100
50
0
*
No Heat Shock
20 min Heat Shock

**Fig. S4. No evidence for induction of the heat shock response in the developing prothoracic gland (PG).** (A) Hoechst staining for DNA (white), immunostaining for Hsp70 (green), and CD8-RFP expression (magenta) in the prothoracic gland of wandering 3rd instar larvae at ~120 h after egg laying (hAEL) in animals with or without a 20 min heat shock. Hsp70 signal was detected only in heat-shocked PGs. Scale bar, 20 μm. (B) Quantitation of relative Hsp70 immunofluorescence in PGs with or without heat shock from (A). n = 6 and 4 PGs for each condition. Statistical significance was determined using Welch's t test. Data are presented as mean ± 95% CI. $P < 0.05$ (*), $P < 0.01$ (**), $P < 0.001$ (***), $P < 0.0001$ (****).

**Fig. S5.**

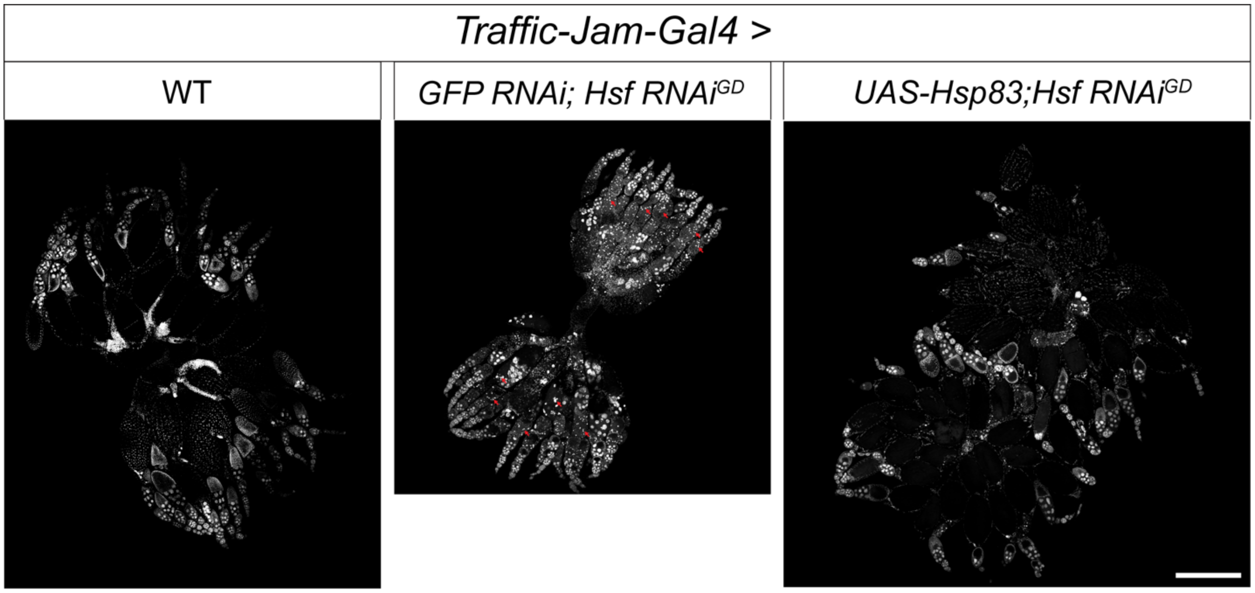

**Fig. S5. Follicle cell-specific *Hsf* knockdown causes follicle degeneration.** Hoechst staining for DNA (white) in the ovary of control and follicle cell-specific knockdown of *Hsf* (*Hsf RNAi)* females, with or without HSP83 re-expression (*UAS-Hsp83*). *Hsf RNAi* resulted in degenerating follicles, as indicated by condensed DNA staining (red arrows). Co-expression of HSP83 suppressed this defect. Scale bar, 500 μm.

### Table S1

| Key Resources Table | | | | |
|---|---|---|---|---|
| **Reagent type (species) or resource** | **Designation** | **Source or reference** | **Identifiers** | **Additional information** |
| genetic reagent (Drosophila melanogaster) | Hsf$^{1}$ | Bloomington Drosophila Stock Center | FLYB:FBst0005491;<br>RRID:BDSC_5491 | Genotype: net$^{1}$ cn$^{1}$ Hsf$^{1}$/CyO<br>Note: Hsf loss-of-function mutant |
| genetic reagent (Drosophila melanogaster) | Hsf$^{03091}$ | Bloomington Drosophila Stock Center | FLYB:FBst0011271;<br>RRID:BDSC_11271 | Genotype: cn1 P{PZ}Hsf$^{03091}$/CyO; ry$^{506}$<br>Note: Hsf loss-of-function mutant |
| genetic reagent (Drosophila melanogaster) | Df(2R)ED3610 | Bloomington Drosophila Stock Center | FLYB:FBst0009066;<br>RRID:BDSC_9066 | Genotype: w$^{1118}$; Df(2R)ED3610, P{3'.RS5+3.3'}ED3610/In(2L)Cy$^{L}$t$^{R}$ In(2R)Cy, Duox$^{Cy}$ amos$^{Roi-1}$<br>Note: Hsf deficiency |
| genetic reagent (Drosophila melanogaster) | Hsf-Halo | Gift from Dr. Carl Wu<br>Tang et al., 2022 | | Genotype: w*; P{Hsf.Halo}attP2<br>Note: Hsf rescue construct |
| genetic reagent (Drosophila melanogaster) | Hsf-RNAi$^{GD}$ | Vienna Drosophila Resource Center | FLYB:FBst0462121;<br>SKU#:GD-37699 | Genotype: w$^{1118}$; P{GD4464}v37699<br>Note: Expresses shRNA for Hsf knock down using RNAi |
| genetic reagent (Drosophila melanogaster) | Hsf-RNAi$^{KK}$ | Vienna Drosophila Resource Center | FLYB:FBst0480645;<br>SKU#:KK-108851 | Genotype: P{KK100723}VIE-260B<br>Note: Expresses shRNA for Hsf knock down using RNAi |
| genetic reagent (Drosophila melanogaster) | phm-Gal4 | Bloomington Drosophila Stock Center | FLYB:FBst0009066;<br>RRID:BDSC_80577 | Genotype: y$^{1}$ w*; P{phtm-GAL4.O}22<br>Note: Expresses GAL4 in prothoracic gland, but not in the corpus allatum. |
| genetic reagent (Drosophila melanogaster) | UAS-CD8-RFP | Gift from Dr. Dan Bergstralh | | Genotype: y$^{1}$ w*; P{UAS-mCD8.mRFP.LG}18a<br>Note: Construct on the second chromosome. Express membrane targeting RFP. Used to label prothroacic gland membrane. |
| genetic reagent (Drosophila melanogaster) | Dref-RNAi | Bloomington Drosophila Stock Center | FLYB:FBst0031941;<br>RRID:BDSC_31941 | Genotype: y$^{1}$ v$^{1}$; P{TRiP.JF02232}attP2<br>Note: Expresses shRNA for Dref knock down using RNAi |
| genetic reagent (Drosophila melanogaster) | Hsp83-RNAi | Vienna Drosophila Resource Center | FLYB:FBst0470805;<br>SKU#:GD-7716 | Genotype: w$^{1118}$; P{GD1202}v7716<br>Note: Expresses shRNA for Hsp83 knock down using RNAi |
| genetic reagent (Drosophila melanogaster) | GFP-RNAi | Bloomington Drosophila Stock Center | FLYB:FBst0009331;<br>RRID:BDSC_9331 | Genotype: w$^{1118}$; P{UAS-GFP.RNAi.R}143<br>Note: Expresses shRNA for GFP knock down using RNAi, used as a control for the total expression level of UAS construct. |
| genetic reagent (Drosophila melanogaster) | Hsp83-GFP | Gift from Dr. Valeria Palumbo<br>Tariq et al., 2009, Palumbo et al., 2020 | | Genotype: Hsp83-GFP<br>Note:The GFP was fused C-terminally to the Hsp83 CDS. Used to assess the expression level of Hsp83. |
| genetic reagent (Drosophila melanogaster) | UAS-Hsp83 | Bloomington Drosophila Stock Center | FLYB:FBst0058469;<br>RRID:BDSC_58469 | Genotype: y$^{1}$ w*; PBac{UAS-Hsp83.Z}VK00037<br>Note: Used to express Hsp83 for rescue experiments |
| genetic reagent (Drosophila melanogaster) | AB1-Gal4 | Bloomington Drosophila Stock Center | FLYB:FBst0001824;<br>RRID:BDSC_1824 | Genotype: y$^{1}$ w*; P{GawB}AB1<br>Note: Expresses GAL4 in salivary gland (basal expression of P{GawB}). |
| genetic reagent (Drosophila melanogaster) | ptc-Gal4 | Bloomington Drosophila Stock Center | FLYB:FBst0002017;<br>RRID:BDSC_2017 | Genotype: w*; P{GawB}ptc$^{559.1}$<br>Note: Expresses GAL4 in salivary gland. |
| genetic reagent (Drosophila melanogaster) | Cg-Gal4 | Bloomington Drosophila Stock Center | FLYB:FBst0007011;<br>RRID:BDSC_7011 | Genotype: w$^{1118}$; P{Cg-GAL4.A}2<br>Note: Expresses GAL4 in hemocytes (also fat body and lymph gland) under the control of a 2.7kb regulatory region located between Cg25C and vkgs. |
| genetic reagent (Drosophila melanogaster) | koi$^{84}$/CyO,GFP | Gift from Dr. Janice A. Fischer<br>Kracklauer et al., 2007 | | Genotype: koi$^{84}$/CyO,GFP<br>Note: Second chromosome balancer with GFP. |
| genetic reagent (Drosophila melanogaster) | UAS-TARDBP.J/CyO,<br>P{2xTb1-RFP}CyO | Bloomington Drosophila Stock Center | FLYB:FBst0051371;<br>RRID:BDSC_51371 | Genotype: w$^{1118}$; P{UAS-TARDBP.J}6M/CyO, P{2xTb$^{1}$-RFP}CyO<br>Note: Second chromosome balancer with RFP (Tb[1]-RFP fusion protein resulting in cuticular fluorescence and a shortened body axis). |
| genetic reagent (Drosophila melanogaster) | MS1096-Gal4 | Bloomington Drosophila Stock Center | FLYB:FBst0008860;<br>RRID:BDSC_8860 | Genotype: w$^{1118}$ P{GawB}Bx$^{MS1096}$<br>Note: Expresses GAL4 in the dorsal wing disc. |
| genetic reagent (Drosophila melanogaster) | traffic-jam-Gal4 | Gift from Dr. Dan Bergstralh | | Genotype: traffic-jam-Gal4/CyO<br>Note: Expresses GAL4 in the follicle cells. |
| Antibody | rabbit anti-HSF | Gift from Dr. Susan Lindquist, origionally from Dr. Tim Westwood (Westwood et al., 1991) | | Working concentration: 1:1000 |

| | | | | |
|---|---|---|---|---|
| Antibody | rabbit anti-Hsp70 | Gift from Dr. Susan Lindquist (Velazquez et al., 1983) | Clone 7FB | Working concentration: 1:1000 |
| Antibody | rabbit anti-Dib | Gift from Dr. Naoki Yamanaka | | Working concentration: 1:1000 |
| Antibody | rabbit anti-HSP90 | Cell Signaling. Cat No. 4874 | RRID:AB_2121214 | Working concentration: 1:1000 |
| Antibody | mouse anti-α-Tubulin | Cell Signaling. Cat No. 3873 | RRID:AB_1904178 | Working concentration: 1:1000 |
| Antibody | Goat anti-rabbit IgG Alexa Fluor 488 | Invitrogen. Cat No. A-11008 | RRID:AB_143165 | Working concentration: 1:10000<br>Note: pre-cleared by overnight incubation at 4 °C with wild-type larval tissues |
| Antibody | Goat anti-rabbit IgG Alexa Fluor 633 | Invitrogen. Cat No. A-21070 | RRID:AB_2535731 | Working concentration: 1:10000<br>Note: pre-cleared by overnight incubation at 4 °C with wild-type larval tissues |
| Antibody | IRDye® 800CW Goat anti-Rabbit IgG | Li-COR. Cat No. 926-32211 | RRID:AB_621843 | Working concentration: 1:10000 |
| Antibody | IRDye® 680RD Goat anti-Mouse IgG | Li-COR. Cat No. 926-68070 | RRID:AB_10956588 | Working concentration: 1:10000 |
| Chemical compound, drug | Hoechst 33342 | Thermo Fisher. Cat No. 62249 | | Dilution 1 to 1:000 |
| Chemical compound, drug | 20-Hydroxyecdysone | Sigma-Aldrich. SKU: H5142-25MG | | |
| Mounting medium | Aqua-Poly/Mount | Polysciences. Cat No. 18606 | | |
| Fly food | Instant Drosophila Medium | Carlolina Biological Supply. Cat No. 173200 | | Used to make Okamoto food (Okamoto et al., 2018) |
| Media component | Agar gelidium | Mooragar. Cat No. 41084 | | Used to make Okamoto food (Okamoto et al., 2018) |
| Media component | Clintose dextrose | Fisher. Cat No. NC178886 | | Used to make Okamoto food (Okamoto et al., 2018) |
| Media component | Quaker enriched cornmeal | Walmart | | Used to make Okamoto food (Okamoto et al., 2018) |
| Media component | Methyl paraben | VWR. Cat No. IC10234105 | | Used to make Okamoto food (Okamoto et al., 2018) |
| Media component | Propionic acid | Fisher. Cat No. AC149300010 | | Used to make Okamoto food (Okamoto et al., 2018) |

### Table S2

| Figure | Panel | Genotype |
|---|---|---|
| Figure 1 | A | *Oregon R* |
| | | *Hsf*$^{1}$*/Hsf*$^{03091}$ |
| | | *Hsf*$^{1}$*/Hsf*$^{03091}$*; Hsf-Halo* |
| | B | *Oregon R* |
| | | *Hsf*$^{1}$*/Hsf*$^{03091}$ |
| | | *Hsf*$^{03091}$*/Df(2R)ED3610* |
| | | *Hsf*$^{1}$*/Df(2R)ED3610* |
| | | *Hsf*$^{1}$*/Hsf*$^{03091}$*; Hsf-Halo* |
| | C | *Oregon R* |
| | | *Hsf*$^{1}$*/Hsf*$^{03091}$ |
| | | *Hsf*$^{1}$*/Hsf*$^{03091}$*; Hsf-Halo* |
| Figure 2 | A, B, and C | *Hsp83-GFP* |
| | | *Hsf*$^{1}$*/CyO, P{2xTb1-RFP}CyO; Hsp83-GFP or Hsf*$^{03091}$*/CyO, P{2xTb1-RFP}CyO; Hsp83-GFP* |
| | | *Hsf*$^{1}$*/Hsf*$^{03091}$*; Hsp83-GFP* |
| | D | *Oregon R* |
| | | *Hsf*$^{1}$*/Hsf*$^{03091}$ |
| Figure 3 | B-F | *UAS-mCD8.mRFP/+; phm-Gal4/+* |
| | | *UAS-mCD8.mRFP/+; phm-Gal4/Hsf RNAi*$^{GD}$ |
| | | *UAS-mCD8.mRFP/Hsf RNAi*$^{KK}$*; phm-Gal4/+* |
| Figure 4 | A-D | *Hsp83-GFP/+; phm-Gal4/+* |
| | | *Hsp83-GFP/+; phm-Gal4/Hsf RNAi*$^{GD}$ |
| | | *Hsp83-GFP/+; phm-Gal4/Dref RNAi* |
| | E-G | *UAS-mCD8.mRFP/+; phm-Gal4/+* |
| | | *UAS-mCD8.mRFP/+; phm-Gal4/Hsp83 RNAi* |
| Figure 5 | A-D | *UAS-mCD8.mRFP/+; phm-Gal4/+* |
| | | *UAS-mCD8.mRFP/GFP RNAi; phm-Gal4/Hsf RNAi*$^{GD}$ |
| | | *UAS-mCD8.mRFP/UAS-Hsp83; phm-Gal4/Hsf RNAi*$^{GD}$ |
| Figure 6 | A, C, and D | *AB1-Gal4/+* |
| | | *AB1-Gal4/+; Hsf RNAi*$^{GD}$*/+* |
| | | *AB1-Gal4/GFP RNAi; Hsf RNAi*$^{GD}$*/+* |
| | | *AB1-Gal4/UAS-Hsp83; Hsf RNAi*$^{GD}$*/+* |
| | B, C, and D | *AB1-Gal4/+* |
| | | *AB1-Gal4/+; Hsp83 RNAi/+* |
| | E-H | *Hsp83-GFP/+; ptc-Gal4/+* |
| | | *Hsp83-GFP/+; ptc-Gal4/Hsf RNAi*$^{GD}$ |
| | | *Hsp83-GFP/+; ptc-Gal4/Dref RNAi* |
| Figure 7 | A and B | *MS1096-Gal4/+* |
| | | *GFP RNAi/+; Hsf RNAi*$^{GD}$*/+; MS1096-Gal4/+* |
| | | *UAS-Hsp83/+; Hsf RNAi*$^{GD}$*/+; MS1096-Gal4/+* |
| | C and D | *Cg-Gal4/+* |
| | | *Cg-Gal4/GFP RNAi; Hsf RNAi*$^{GD}$*/+* |
| | | *Cg-Gal4/UAS-Hsp83; Hsf RNAi*$^{GD}$*/+* |
| Supplementary Figure 1 | A | *UAS-mCD8.mRFP/+; phm-Gal4/+* |
| | | *UAS-mCD8.mRFP/+; phm-Gal4/Hsf RNAi*$^{GD}$ |
| | B | *UAS-mCD8.mRFP/+; phm-Gal4/+* |
| | | *UAS-mCD8.mRFP/+; phm-Gal4/Hsf RNAi*$^{GD}$ |
| | | *UAS-mCD8.mRFP/Hsf RNAi*$^{KK}$*; phm-Gal4/+* |
| | C and D | *UAS-mCD8.mRFP/+; phm-Gal4/+* |
| | | *UAS-mCD8.mRFP/Hsf RNAi*$^{KK}$*; phm-Gal4/+* |
| Supplementary Figure 2 | A | *Hsp83-GFP/+; phm-Gal4/+* |
| | | *Hsp83-GFP/+; phm-Gal4/Dref RNAi* |
| | B, D, and E | *UAS-mCD8.mRFP/+; phm-Gal4/+* |
| | | *UAS-mCD8.mRFP/+; phm-Gal4/Hsp83 RNAi* |
| | C | *UAS-mCD8.mRFP/+; phm-Gal4/Hsp83 RNAi* |
| Supplementary Figure 3 | A-D | *UAS-mCD8.mRFP/+; phm-Gal4/+* |
| | | *UAS-mCD8.mRFP/UAS-Hsp83; phm-Gal4/+* |
| Supplementary Figure 4 | A and B | *UAS-mCD8.mRFP; phm-Gal4* |
| Supplementary Figure 5 | | *Traffic-Jam-Gal4/+* |
| | | *Traffic-Jam-Gal4/GFP RNAi; Hsf RNAi*$^{GD}$*/+* |
| | | *Traffic-Jam-Gal4/UAS-Hsp83; Hsf RNAi*$^{GD}$*/+* |
